# Predicting cancer prognosis and drug response from the tumor microbiome

**DOI:** 10.1101/2020.07.21.214148

**Authors:** Leandro C. Hermida, E. Michael Gertz, Eytan Ruppin

## Abstract

Tumor gene expression is predictive of patient prognosis in some cancers. However, RNA-seq and whole genome sequencing data contain not only reads from host tumor and normal tissue, but also reads from the tumor microbiome, which can be used to infer the microbial abundances in each tumor. Here, we show that tumor microbial abundances, alone or in combination with tumor gene expression data, can predict cancer prognosis and drug response to some extent – microbial abundances are significantly less predictive of prognosis than gene expression, although remarkably, similarly as predictive of drug response, but in mostly different cancer-drug combinations. Thus, it appears possible to leverage existing sequencing technology, or develop new protocols, to obtain more non-redundant information about prognosis and drug response from RNA-seq and whole genome sequencing experiments than could be obtained from tumor gene expression or genomic data alone.

## Introduction

The Cancer Genome Atlas (TCGA), available from the NCI Genomic Data Commons (GDC)^1^, provides RNA-seq and whole genomic sequencing (WGS) data for thousands of cases across dozens of cancer types. RNA-seq data is typically used to measure the expression of human genes, and there is a long history linking tumor gene expression to cancer outcomes^2–8^. Milanez-Almeida et al.^9^ recently showed that gene expression from TCGA RNA-seq data could predict overall survival (OS) or progression-free interval (PFI) better than classical clinical prognostic covariates – age at diagnosis, gender, and tumor stage. Importantly, Milanez-Almeida et al. used a data-driven machine learning (ML) based approach which selected features that were predictive of and correlated with prognosis, rather than approaches based on classical statistics or biological knowledge that chose features *a priori*.

Research into the human tumor microbiome has been rapidly expanding, and multiple laboratories have attempted to utilize existing technologies and data to identify microbes and quantify their abundance within human tumors compared to adjacent normal tissue. RNA-seq and WGS data not only contain human sequencing reads, but also reads from the local intratumor microbiome that are typically filtered out from the data when analyzing human gene expression or genomic alterations. Poore et al.^10^ recently developed a computational workflow, using two orthogonal microbial detection pipelines, to estimate, decontaminate, normalize, and batch effect correct microbial abundances from human high-throughput sequencing data. They applied this workflow to create a first-of-its-kind comprehensive dataset of pan-cancer tumor microbial abundances derived from WGS or RNA-seq data for the entire TCGA cohort.

Our central research questions then were, 1) does a data-driven ML approach reveal that tumor microbial abundances in TCGA data, quantified from these reads, are predictive of cancer prognosis or drug response, 2) what microbial genera are potentially predictive biomarkers of prognosis or drug response, 3) how do these models compare to equivalent models based on tumor gene expression data, and 4) does combining both microbial abundance and gene expression features produce models and select combinations of genes and microbial genera that are more predictive of prognosis or drug response than models from each individual data type? We used the processed microbial abundances directly from the Poore et al. dataset to build predictive models of prognosis and drug response for TCGA. We also used TCGA RNA-seq read counts to build equivalent predictive models for comparison. As a positive control, we also showed that our prognosis ML modeling methods, which differed somewhat from Milanez-Almeida et al.^9^, identified a similar set cancers and outcomes for which gene expression was predictive of prognosis.

## Results

### Tumor microbial abundances are substantially less predictive of prognosis than gene expression

An overview of the analytical workflow is presented in **Fig. 1**. It has four major parts, 1) data download and preprocessing, 2) prognosis and drug response ML modeling, 3) model evaluation and scoring, and 4) further feature analysis. A more detailed technical description of the analysis pipeline and computational methods is provided in Methods.

**Figure 1.**
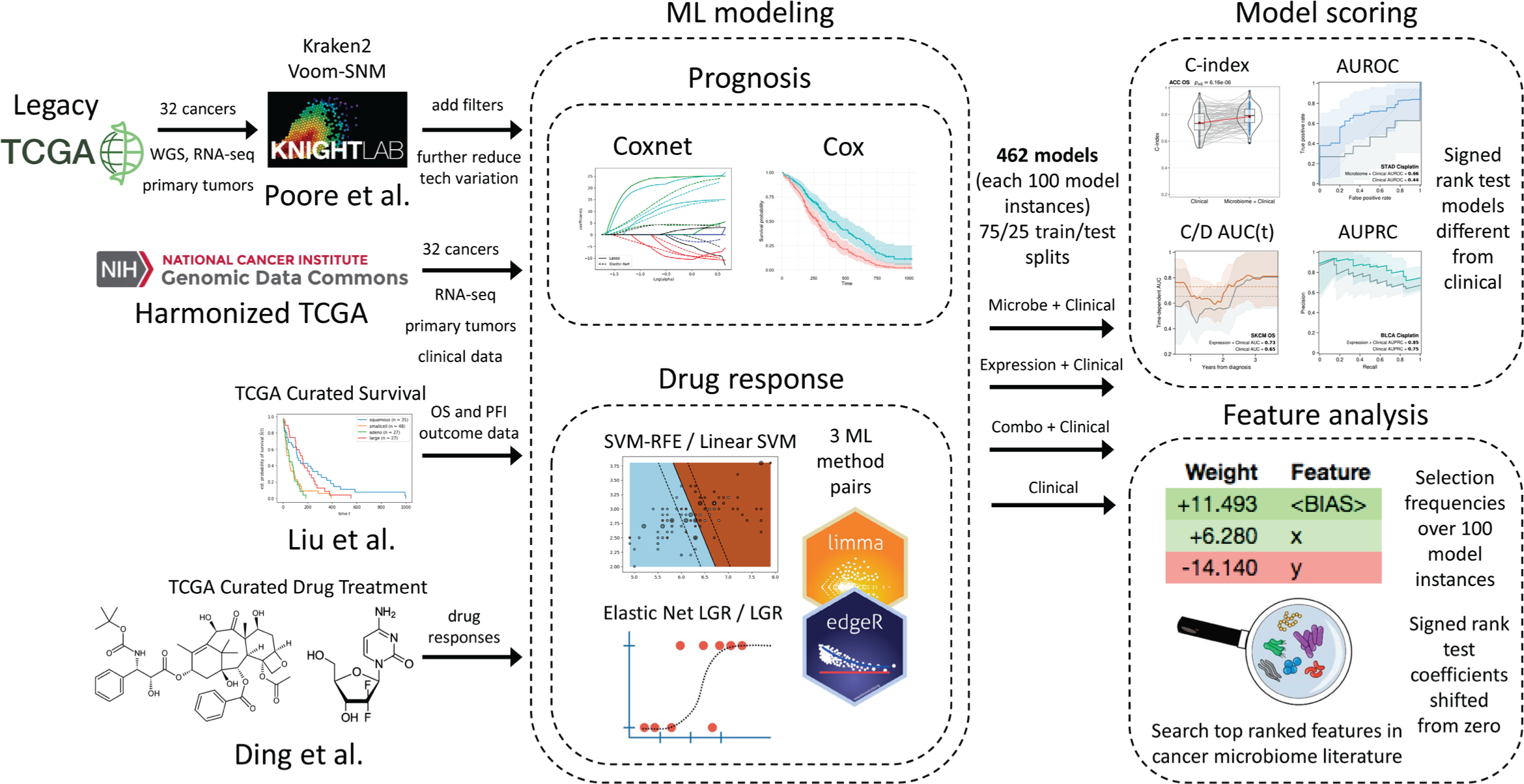
Analysis pipeline overview. Download and data preprocessing (**left**) of Poore et al. TCGA primary tumor Kraken2 Voom-SNM microbial abundances with additional filters to reduce technical variation, NCI Genomic Data Commons (GDC) harmonized TCGA primary tumor RNA-seq counts and clinical data, Liu et al. TCGA curated overall survival (OS) and progression-free interval (PFI) outcome data, and Ding et al. TCGA curated drug response clinical data. Prognosis machine learning (ML) modeling (**middle**) of microbial abundance, gene expression, and combined data types with clinical covariates for each cancer using penalized Cox with elastic net penalties (Coxnet) against matched clinical covariate-only models using standard Cox regression. Drug response classification ML modeling of the same data types with clinical covariates for each cancer-drug combination using three ML approaches, 1) SVM-RFE, elastic net logistic regression (LGR), and limma-trend (microbial and combined data types) or edgeR (gene expression) differential analysis feature scoring and selection with L2 penalized LGR. Matched clinical covariate-only modeling performed with L2 penalized linear SVM or LGR. ML modeling generates 100 model instances for each model from 75/25 train/test randomly shuffled and stratified dataset splits. ML model instance scoring (**right top**) using concordance index (C-index) and time-dependent cumulative/dynamic AUC (C/D AUC(t)) for prognosis models and area under receiver-operating characteristic curve (AUROC) and area under the precision-recall curve (AUPRC) for drug response models. Significance of model performance improvement over matched clinical covariate-only model determined by signed rank test of C-index or AUROC scores between each matched model instance for prognosis and drug response models, respectively. Feature analysis (**right bottom**) performed using model instance coefficients and selection frequencies. Overall feature importance ranking and significance determined by signed rank test of model instance feature coefficients shifting from zero and filtering of top features for selection frequency ≥ 20%.

We built OS and PFI gene expression ML models of 32 TCGA tumor types (see **Supplementary Data 1** for cohort information) using the Coxnet^11^ algorithm, which jointly selects the most predictive subset of features via cross-validation (CV) while simultaneously being able to control for prognostic clinical covariates. In our models, we included and controlled for the clinical covariates age at diagnosis, gender, and tumor stage. For comparison, we also built standard Cox regression models based on the clinical covariates alone. We evaluated the predictive performance of our models using Harrell’s concordance index (C-index), which is a metric of survival model predictive accuracy. Each model analysis generated 100 model instances and C-index scores from randomly shuffled train-test CV splits on the data. We found 33 OS and PFI models for 21 tumor types that had a mean C-index score ≥ 0.6 and significantly outperformed their corresponding clinical covariate-only models (**Fig. 2a & c, Supplementary Figs. 1a, 2a**). Our models were predictive of prognosis in 11 of the same 13 tumor types that were reported by Milanez-Almeida et al.^9^ (**Supplementary Table 1**). We did not analyze one tumor type that Milanez-Almeida did, acute myeloid leukemia (LAML), because Poore et al. excluded it from their analysis. Among the cancers and outcomes that Milanez-Almeida et al. analyzed, our methodology produced predictive models for four additional tumor types: breast cancer (BRCA), cervical squamous cell carcinoma (CESC), sarcoma (SARC), and uterine corpus endometrial carcinoma (UCEC), as well as quite a few predictive models for additional cancers and outcomes that were not analyzed in their study (**Supplementary Table 1**). We also evaluated prognosis model performance by calculating the time-dependent, cumulative/dynamic area under the curve (AUC^C/D^(t))^12, 13^, which is an extension of the area under the receiver-operating characteristic curve (AUROC) for continuous outcomes and can provide a more detailed resolution picture of predictive performance throughout the test outcome time range compared to the C-index score. Although 33 of our OS and PFI gene expression models had a statistically significant C-index score improvement compared to clinical covariates alone, only 22 of these models showed an improvement in AUC^C/D^(t), where the improvement in mean AUC^C/D^(t) over the entire test time range after diagnosis was ≥ 0.025 (**Supplementary Figs. 1b, 2b**).

**Figure 2.**
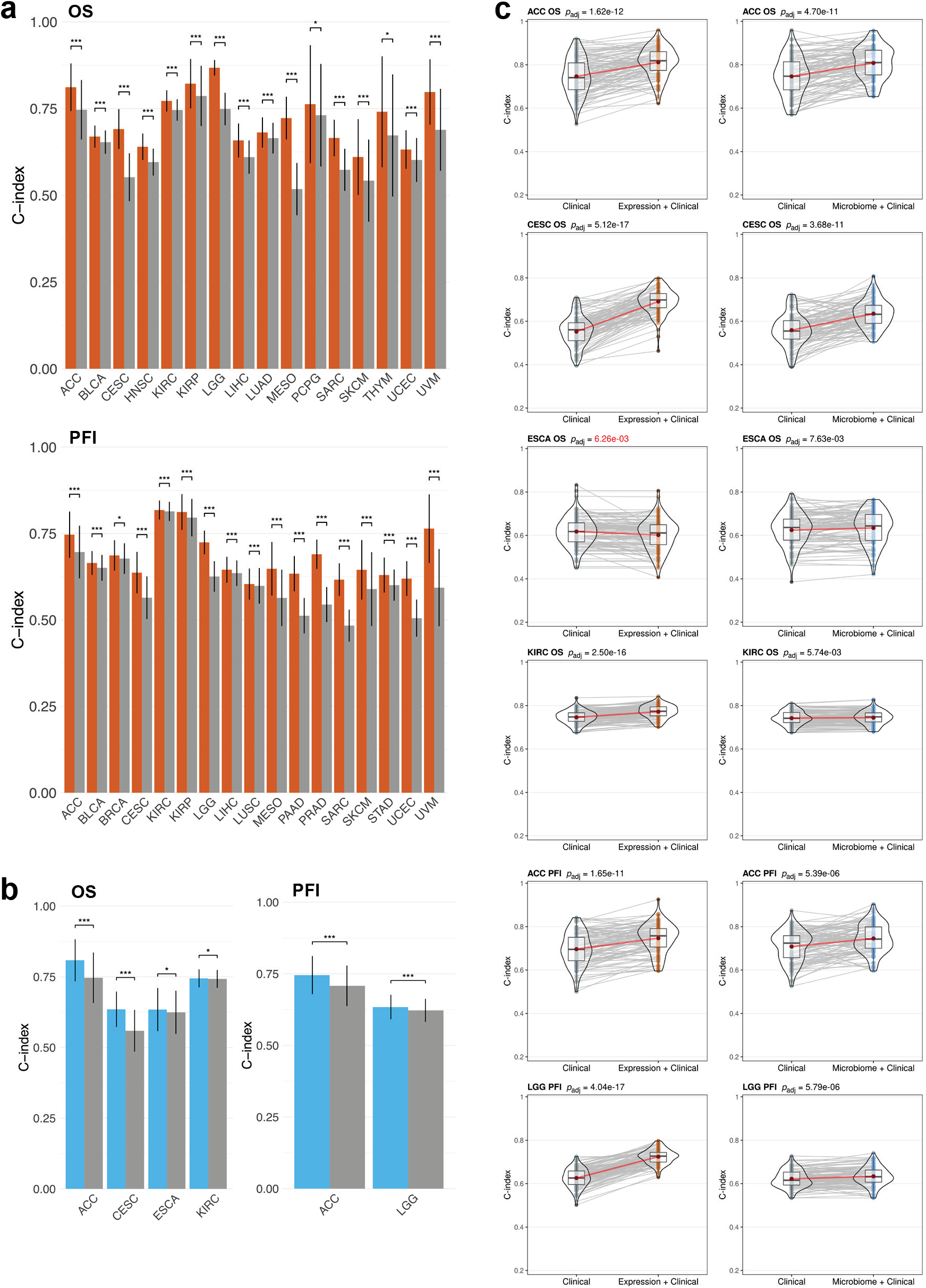
Performance of gene expression and microbial abundance prognosis prediction models where features add predictive power to clinical covariates. Mean C-index scores for **(a)** gene expression with clinical covariate models (orange) and **(b)** microbial abundance with clinical covariate models (blue) vs clinical covariate- only models (grey). Error bars denote standard deviations. Significance: * ≤ 0.01, ** ≤ 0.001, *** ≤ 0.0001. **(c)** C-index score violin density plots for the six models where microbial abundance with clinical covariate features outperform clinical covariate-only models. Corresponding gene expression models shown for comparison. Lines connecting points (light grey) represent score pairs from same train-test split on the data. Mean C-index scores shown as red dots with red lines connecting the means. Significance for the prediction improvement over clinical covariate-only models was calculated using a two-sided Wilcoxon signed-rank test and adjusted for multiple testing using the Benjamini-Hochberg method with adjusted p-values shown at top. Adjusted p-values colored in red signify difference where clinical covariate-only model is better.

We applied Coxnet^11^ using the same methodology to build prognosis models using the microbial abundance estimates provided by Poore et al.^10^. We found six microbial abundance models that had a mean C-index score ≥ 0.6 and significantly outperformed their corresponding clinical covariate-only models (**Fig. 2b & c, Supplementary Fig. 3a**). We found that in only two of the six models, microbial abundances outperformed clinical covariates alone in terms of AUC^C/D^(t), where the improvement in mean AUC^C/D^(t) over the entire test time range after diagnosis was ≥ 0.025 (**Supplementary Fig. 3b**). In adrenocortical carcinoma (ACC), microbial features predicted OS significantly better than clinical prognostic covariates starting at approximately 6 years after diagnosis. In CESC, microbial abundances predicted OS better than clinical covariates from approximately 6 months to 10 years after diagnosis. Overall, we found that tumor microbial abundances from Poore et al. were only marginally predictive of prognosis across the TCGA cohort, and that gene expression was a significantly more powerful predictor of prognosis (**Fig. 2, Supplementary Figs. 1-3**).

### Tumor microbial abundances are predictive of chemotherapy drug response in some cancers and in mostly different cancer-drug combinations than gene expression

We next asked whether tumor microbial abundances from pre-treatment biopsies could predict drug response better than the clinical covariates age at diagnosis, gender, and tumor stage alone. TCGA drug response clinical data were obtained from Ding et al.^14^ as described in Methods. Cases with complete response (CR) or partial response (PR) were labeled as responders and those with stable disease (SD) or progressive disease (PD) as non-responders. Thirty TCGA cancer-drug combinations met our minimum dataset size thresholds (see **Supplementary Data 1** for cohort information). We built drug response models using three different ML methods: 1) a variant of the linear support vector machine recursive feature elimination (SVM-RFE) algorithm^15^ that we developed, 2) logistic regression (LGR) with elastic net^16^ (L1 + L2) penalties and embedded feature selection, and 3) logistic regression with an L2 penalty and limma^17^ (for microbial abundance and combined data type datasets) or edgeR^18, 19^ (for RNA-seq count datasets) differential abundance/expression feature scoring and wrapper selection methods (see Methods for details). All three ML methods unconditionally included the clinical covariates – age at diagnosis, gender, and tumor stage – in the model (bypassing feature selection) while selecting the most predictive subset of microbial abundance or gene expression features. For comparison, we built standard linear SVM or LGR models using the clinical covariates alone. We evaluated the predictive performance of drug response models using AUROC. Each analysis generated 100 model instances, AUROC, and area under the precision-recall curve (AUPRC) scores from randomly shuffled train-test CV splits on the data.

We found five microbial abundance cancer-drug combinations that had a mean AUROC score ≥ 0.6 and performed better than clinical covariates alone in at least two out of three ML methods (**Fig. 3**). Three of these cancer-drug combinations involved stomach adenocarcinoma (STAD). We performed the same drug response modeling using TCGA gene expression data and here we found six cancer-drug response combinations that had a mean AUROC score ≥ 0.6 and significantly outperformed their corresponding clinical covariate-only models in at least two out of three ML methods (**Fig. 4**). Only one cancer-drug combination, SARC docetaxel, overlapped between the microbial abundance and gene expression drug response model results, suggesting that tumor microbial abundances have independent predictive power. Even though one of our thresholds for a significant drug response model was a mean AUROC score ≥ 0.6, the 11 total significant models that we found from both data types each had a mean AUROC > 0.7. We also found there was considerable overlap in the selected microbial abundance and gene expression features reported by each ML method (**Fig. 5a & c**) and frequently found a significant correlation between the feature importance rankings reported by each ML method when comparing the two most significant methods in each cancer-drug combination (**Fig. 5b & d**). These results suggest that our significant drug response models and their inferred important features are not the result a specific ML modeling methodology. Overall, our results support the notion that the tumor microbiome may contain information that is predictive of drug response in some cancers, consistent with recent reports^20, 21^.

**Figure 3.**
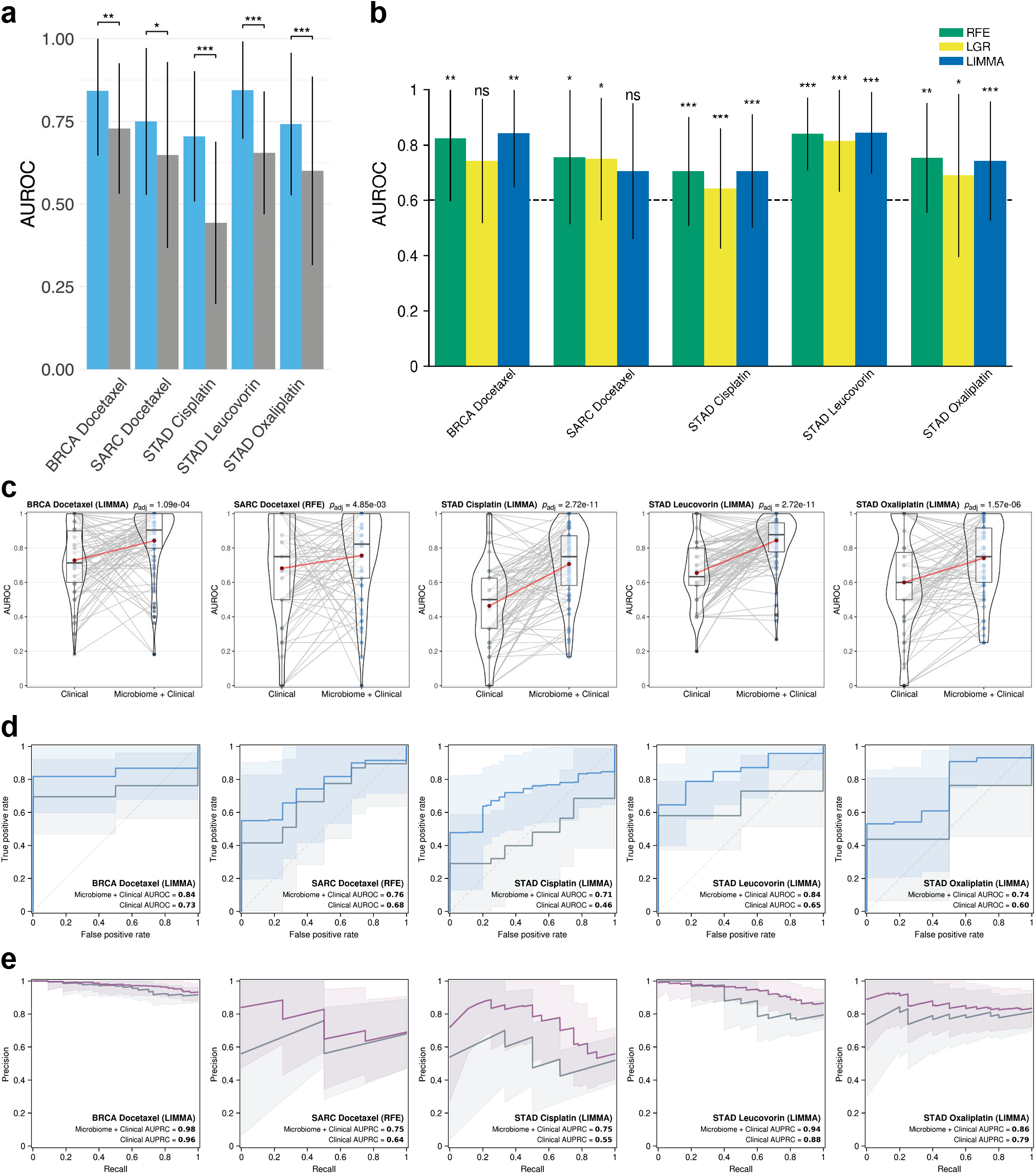
Performance of microbial abundance drug response prediction models in the five cancer-drug combinations where models performed better than clinical covariates alone. **(a)** Mean AUROC scores for microbial abundance with clinical covariate models (blue) vs clinical covariate-only models (grey) and **(b)** mean AUROC scores for each ML method. In both **(a)** and **(b)** error bars denote standard deviations. Significance: * ≤ 0.01, ** ≤ 0.001, *** ≤ 0.0001. **(c)** Violin density plots of AUROC scores for microbial abundance with clinical covariate models vs clinical covariate-only models. Lines connecting points (light grey) represent score pairs from same train-test split on the data. Mean AUROC scores are shown as red dots connected by red lines. **(d)** Mean ROC (blue) and **(e)** precision-recall (PR) curves (purple) for microbial abundance with clinical covariate models vs clinical covariate-only models (grey). Mean AUROC and AUPRC scores shown in legends and shaded areas denote standard deviations. Significance for the prediction improvement over clinical covariate-only models was calculated using a two-sided Wilcoxon signed-rank test and adjusted for multiple testing using the Benjamini-Hochberg method with adjusted p-values shown at top of violin plots in **(c)**. In **(c-e)** results for the modeling method that had the most significant Wilcoxon signed-rank test are shown.

**Figure 4.**
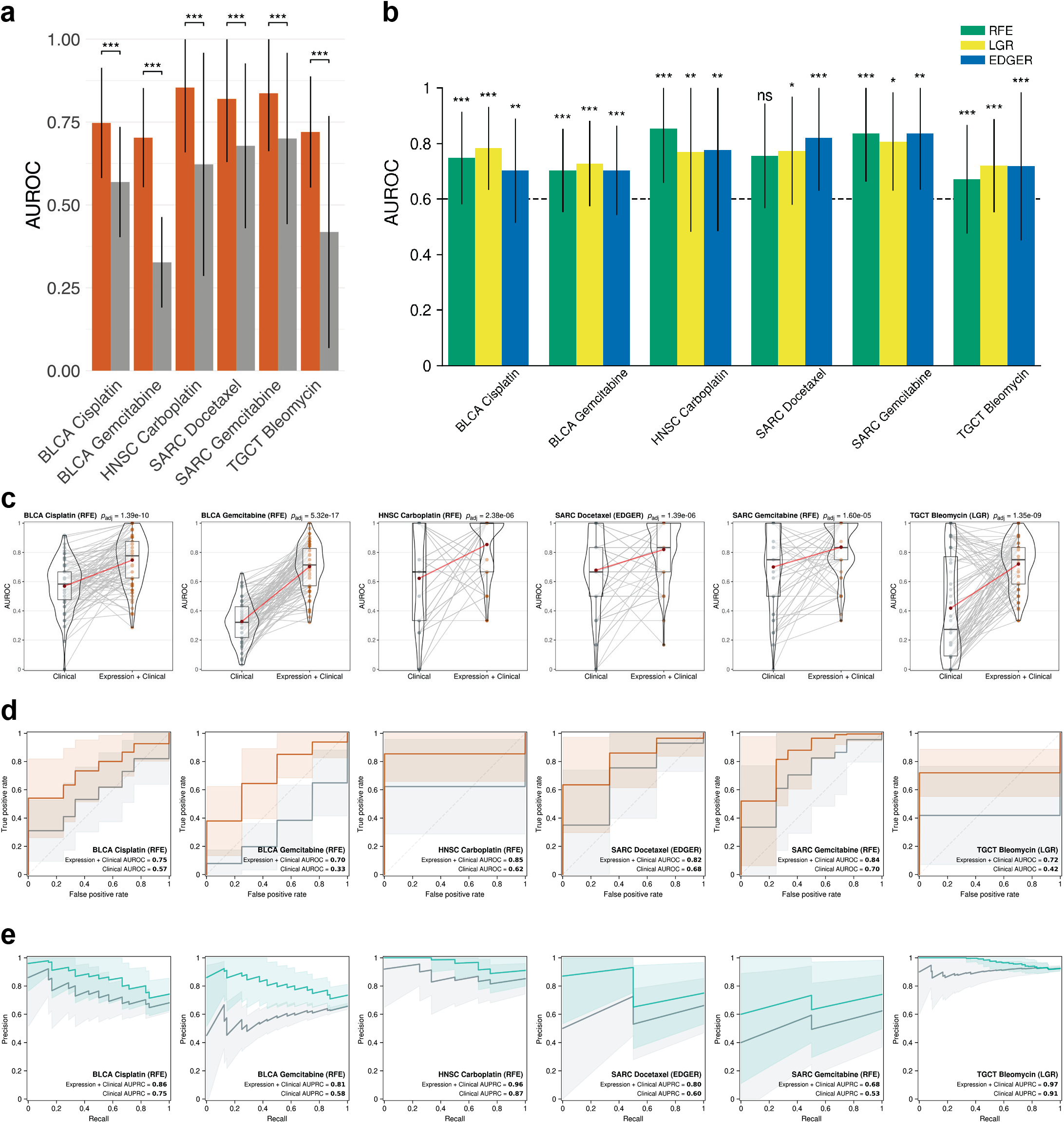
Performance of gene expression drug response prediction models in the six cancer-drug combinations where models performed better than clinical covariates alone. **(a)** Mean AUROC scores for gene expression with clinical covariate models (orange) vs clinical covariate-only models (grey) and **(b)** mean AUROC scores for each ML method. In both **(a)** and **(b)** error bars denote standard deviations. Significance: * ≤ 0.01, ** ≤ 0.001, *** ≤ 0.0001. **(c)** Violin density plots of AUROC scores for gene expression with clinical covariate models vs clinical covariate-only models. Lines connecting points (light grey) represent score pairs from same train-test split on the data. Mean AUROC scores are shown as red dots connected by red lines. **(d)** Mean ROC (orange) and **(e)** precision-recall (PR) curves (green) for gene expression with clinical covariate models vs clinical covariate-only models (grey). Mean AUROC and AUPRC scores shown in legends and shaded areas denote standard deviations. Significance for the prediction improvement over clinical covariate-only models was calculated using a two-sided Wilcoxon signed-rank test and adjusted for multiple testing using the Benjamini-Hochberg method with adjusted p-values shown at top of violin plots in **(c)**. In **(c-e)** results for the modeling method that had the most significant Wilcoxon signed-rank test are shown.

**Figure 5.**
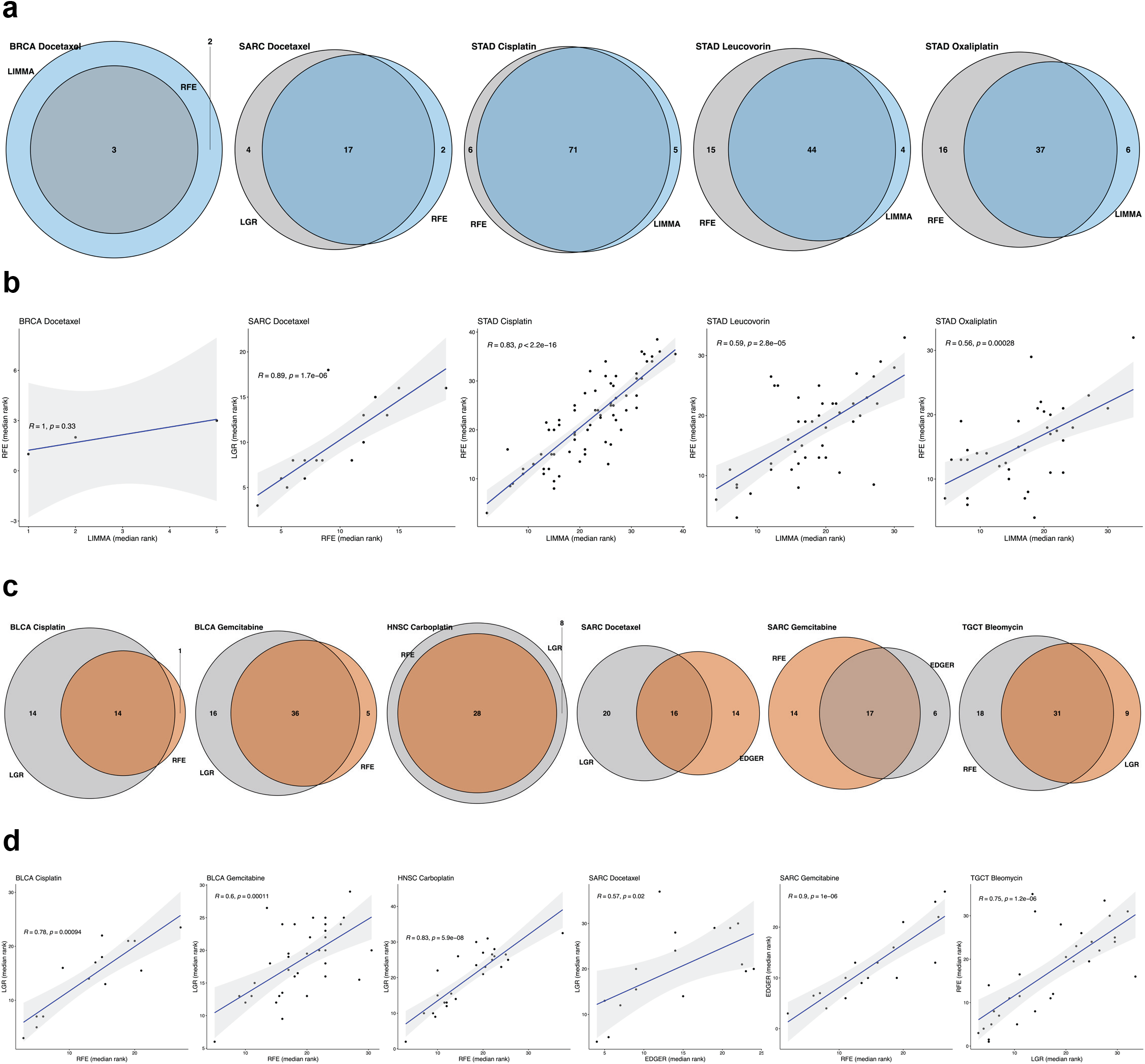
Comparison of drug response model top-ranked selected features by each ML method. For each drug response model, we selected the two best ML methods by significance for the prediction improvement over their respective clinical covariate-only model. **(a, c)** Venn diagrams for microbial abundance **(a)** or gene expression **(c)** models comparing the number of features individually selected by each ML method, and the intersection of the two ML methods. **(b, d)** Spearman rank correlation plots for microbial abundance **(b)** or gene expression **(d)** models showing that the median rank of features (among the 100 model instances in which the feature was selected) often correlated between the two most significant ML methods. The best method is shown on the x-axis, the second best on the y-axis.

### Combining tumor microbial abundance and gene expression features adds a modest predictive improvement in some cancers

Finally, we investigated if models built from combining microbial abundance and gene expression features would result in an improvement in predictive power over their corresponding single data type models. Combining data types resulted in a modest predictive improvement in only three prognosis models: SARC OS, STAD PFI, and thymoma (THYM) OS (**Supplementary Fig. 4a**). Although this improvement was not statistically significant in terms of C-index score, the AUC^C/D^(t) metric showed a clear improvement in prognostic predictive power for these models, where the improvement in mean AUC^C/D^(t) over the entire time range after diagnosis was ≥ 0.025 compared to their respective single data type models. We also found five combined data type drug response models which performed significantly better than clinical covariates alone, although none of these models reached statistical significance when compared to their respective single data type models in terms of improvement in AUROC score, but one of these models, for BLCA cisplatin, did show an improvement in AUROC ≥ 0.025 compared to its corresponding single data type models (**Supplementary Fig. 4b-c**).

### Evaluating the robustness of drug response models

Some of the TCGA drug response cohorts used in our study were of limited size and this could have an impact on the robustness of our analysis (see **Supplementary Data 1** for cohort size information). To study this issue further, we evaluated the significance of model scores using a class label permutation test. We shuffled dataset class labels 1000 times and each time ran the outer CV procedure on the permuted dataset, where for each CV iteration we fit a model instance and calculated an AUROC score. We then calculated a p-value from the fraction of permuted scores that were greater than or equal to the true score. Three of the five microbial abundance drug response models that were reported above to have perform significantly better than clinical covariates alone had a permutation test p-value < 0.05 and the remaining two, for stomach adenocarcinoma (STAD) cisplatin and oxaliplatin, had p-values < 0.08 (**Fig. 6a**). Permutation test scores and significance for microbial abundance models were similar regardless of the modeling method used (**Supplementary Fig. 5a**). Five of the six gene expression drug response models that performed significantly better than clinical covariates alone had a permutation test p-value < 0.05 (**Fig. 6c**). Again here, permutation test scores and significance were similar regardless of the modeling method used (**Supplementary Fig. 6a**). The testicular germ cell tumor (TGCT) bleomycin gene expression model did not quite reach significance, though it is worth mentioning that for the edgeR feature selection and L2 logistic regression modeling method it was close (p = 0.077).

**Figure 6.**
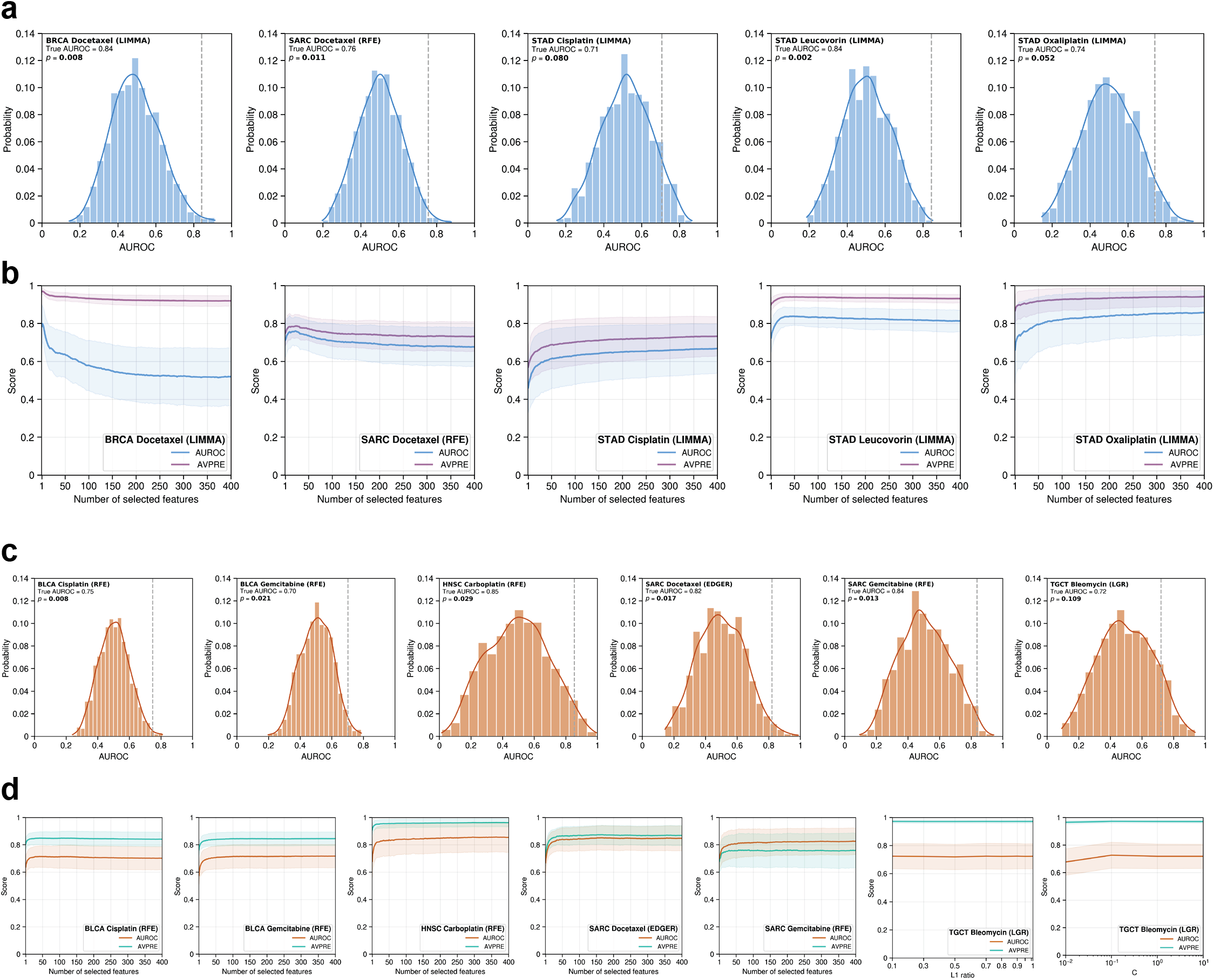
Evaluation of drug response model robustness. Model significance and robustness was further evaluated using a class label permutation test and examination of the effect feature selection had on model performance. Results for the modeling method which had the most significant Wilcoxon signed-rank test are shown. **(a, c)** Permutation test result histograms and significance for microbial abundance **(a)** or gene expression **(c)** models showing the distribution of permutation mean AUROC scores. True mean AUROC score shown as dotted vertical grey line and kernel density estimate shown as a curve over the histogram. **(b, d)** Curves showing the effect that model hyperparameters which control the number of selected features had on mean AUROC and average precision (AVPRE) scores during hyperparameter grid search across all 100 model instances for microbial abundance **(b)** or gene expression **(d)** models. Shaded areas denote standard deviations.

We further evaluated the robustness of our significant drug response models by examining the effect that the number of selected features had on model performance. During the hyperparameter grid search and tuning that occurred in the nested inner CV during each model instance fitting, scores for every combination of hyperparameter setting and inner CV train/validation fold were saved (see Methods for full details). We plotted how these scores were affected by the hyperparameters that controlled feature selection. Our decision to conservatively limit the feature selection search space in our drug response models to a maximum of 400 best scoring features, to reduce model complexity and the possibility of overfitting, appeared sufficient, as scores for our significant models reached a maximum or leveled off well within this search range (**Fig. 6b & d**). In the five microbial abundance models, predictive power was driven by a small number of features in three models, where selecting more features did not contribute to additional predictive power or it added noise (**Fig. 6b**). Even in the remaining two models, most of the predictive power was driven by the top 50 to 100 features. In the six gene expression models, this finding was even more stark, where all the predictive power was achieved by a small number of features in each model (**Fig. 6d**). In all the significant models from both data types, the variance in scores was not significantly affected by the number of selected features and feature-to-sample ratio within our chosen hyperparameter search range. As with the permutation test results, we found the effect that the number of selected features had on model performance was similar regardless of the feature selection or modeling method used (**Supplementary Fig. 5b, 6b**). In summary, these two comprehensive analyses suggest that the significant cancer-drug response combinations found in this study and the most important features inferred from their models represent a potentially real and robust biological signal.

### Feature analysis reveals a wide range of predictive microbial genera

To learn more about the most predictive features, we determined the top microbial genera and top genes (**Supplementary Data 2**) selected by each of the significantly predictive microbial abundance and gene expression models, respectively, according to their selection frequency and model coefficients across the 100 model instances from each analysis. There were 428 distinct microbial genera appearing in at least one prognosis or drug response model. Of these 428 genera, 160 were individually significantly predictive of prognosis or drug response by a Wilcoxon test, indicating that the other genera were significantly predictive in combination. The median number of genera selected per model was 52, with a minimum of 3 (BRCA docetaxel) and a maximum of 78 (STAD cisplatin). Of the 428 genera, 95 were selected in more than one model and only 13 were selected in more than two models. This is consistent with the observation of Nejman et al.^22^ that the tumor microbiome is tumor type specific. The predictive genera we found span all non-eukaryotic domains of life, in total encompassing 365 bacterial, 17 archaeal, and 46 viral genera (**Supplementary Data 2**).

## Discussion

In summary, we find that the microbial abundance estimates generated by Poore et al.^10^ are predictive of cancer patient prognosis and response to chemotherapy in a subset of tumor types, survival outcomes, and treatments. Machine learning methods, such as those applied in this study, are not able to infer causality, but only inform on the positive or negative predictive associations covariates have with the response variable. The potential causal role that those covariates may play in determining patient prognosis or drug response can only be ascertained via dedicated mechanistic studies. Overall, in terms of the number of significant models, based on their cross-validated C-index or AUROC scores and improvement over clinical covariates alone, the tumor microbiome is considerably less predictive than the tumor human transcriptome at predicting patient prognosis, but notably, performs similarly to gene expression at predicting chemotherapy response and in mostly different cancer-drug combinations. Our investigation motivates future studies investigating the role of the tumor microbiome in predicting the response to targeted therapies and immunotherapies.

There are also some limitations to our current study. As we described previously, some TCGA drug response cohorts were of limited size or had relatively few responder or non-responder cases within these cohorts and this could have an impact on the interpretability of the results. Vabalas et al.^23^ conducted a literature review of ML algorithm validation of high-dimensional biological data models with limited sample size and performed their own independent simulation analyses evaluating different techniques. They found that, consistent with previous literature, nested CV was the optimal validation method and gives unbiased performance estimates regardless of sample size. They also found that performing feature selection and other model development steps (e.g., normalization, outlier removal) fully within the inner nested CV is essential to avoid overfitting and to produce unbiased results, and that hyperparameter tuning should ideally also be performed in nested fashion. Finally, they found that performing an adequate number of CV folds was important to reduce bias. Our analyses have followed their observations and recommendations, employing them at every level of model development and evaluation, including additional techniques not reviewed in their work (see Methods for full details).

There are further limitations to this study inherited from limitations in the data, originally raised by Poore et al.^10^ First, the study was retrospective, using existing data from the TCGA. As such, it did not involve any specific protocols to capture microbial reads or to control for contamination. Second, decontamination of such retrospective data is a highly involved and dataset-specific process, which they made great effort to validate. Poore et al. conclude from this validation that the retrospective study of TCGA was successful, and that similar retrospective studies would be valuable. A third point, which they touch on briefly, is that the protocols that were used have limitations with respect to capturing microbial reads and cannot distinguish if the source of microbial reads is intracellular or extracellular, or alive or dead when the sample was taken. Poore et al. suggest, correctly we believe, that additional protocols need to be developed for prospective studies.

Accepting the limitations of the study, we observed certain trends. Proteobacteria and Firmicutes were the most frequent phyla identified as predictive features (**Supplementary Data 2**), followed by Actinobacteria and Bacteroidetes. Among viruses, Herpesvirales were the most frequent. More microbial genera were negatively predictive of drug response or prognosis than those positively predictive (negative for 306/537 features; two-sided binomial test p-value = 0.0014). Firmicutes reversed this trend, being more often positively predictive (positive for 49/82 features, two-sided Fisher’s exact test p-value = 0. 0.0036; **Supplementary Table 2**).

Further examining the predictive features of our significant models and their cancer types, we found that several genera of Firmicutes were predictive of OS in CESC, including genera of *Lactobacillales* were found to be negatively predictive of survival. We also found that the genus *Chlamydia* had an even stronger negatively predictive association with OS in CESC. Notably, though CESC is known to often arise from HPV infection, the presence of other microbial species, in particular the genera *Chlamydia* and *Lactobacillales*, have been reported in the literature to be associated with the risk of developing CESC^24, 25^.

Our prognosis analysis results were different than two recent reports^26, 27^ which found some intratumor microbes that were potentially correlated with prognosis in three TCGA cancers. We did not find that the Poore et al. tumor microbial abundances estimated from TCGA were predictive of OS or PFI in these three cancers using our data-driven, regularized ML computational approach. A few important possible reasons for this difference in results are that different source data and methods were used to perform prognosis analysis compared to these studies. Gnanasekar et al.^26^ analyzed the THCA cohort by tumor subtype, they used harmonized and normalized GDC TCGA data instead of legacy TCGA followed by normalization and batch effect correction as in Poore et al., they only used RNA-seq data instead of WGS and RNA-seq data, they applied different methods for extraction of microbial reads and decontamination, and finally they did not perform any direct analysis of correlation of their derived microbial abundances with survival outcomes. Dohlman et al.^27^ analyzed colorectal cancers (colon (COAD) and rectum (READ) adenocarcinomas) also using harmonized and normalized GDC TCGA data, they used WGS and whole exome sequencing (WXS) data instead of WGS and RNA-seq, they also used different methods for extraction and decontamination of microbial reads, and finally they also applied classical univariate statistics on their entire data to infer correlation with overall survival (OS). While we believe the use of harmonized GDC TCGA data is superior to legacy TCGA, Poore et al. applied robust computational methods to remove technical variation from legacy TCGA data and validated that their approach was effective. We also applied additional filters of TCGA samples to further remove technical variation. We also believe that, in general, applying classical univariate statistics on the entire data to find correlations has the potential to overfit the specific dataset and it does not consider the multivariate nature of high-dimensional biological data like intratumor microbial abundances. A data-centric, multivariate, and regularized ML approach focused on fitting models on training data and evaluating on unseen test data has potential to generalize better and discover whether features are potentially predictive of and correlated with the response variable, such as survival outcomes or drug response.

Looking at our drug response model results, in STAD, tumor microbial abundances were predictive of response to three different drugs: cisplatin, leucovorin, and oxaliplatin. The genus *Helicobacter* was a quantified microbial abundance feature in the Poore et al. dataset although notably, even though it is well established that patients infected with *H. pylori* have an increased risk of developing gastric cancer^28^, *Helicobacter* was not identified as a predictive feature of drug response in our STAD models. This finding is in line with recent research indicating reduced microbial diversity, decreased abundance of *H. pylori*, and enrichment of other mostly commensal bacterial genera in gastric carcinoma^29^. Instead, in STAD we found that known opportunistic bacteria *Cedecea* and *Sphingobacterium* were both strongly negatively predictive of leucovorin response, *Sphingobacterium* was strongly negatively predictive of cisplatin response, and the opportunistic bacteria *Rouxiella* was strongly negative predictive of oxaliplatin response. *Cedecea* and *Sphingobacterium* have been implicated in bacteremia in immunocompromised individuals in rare cases, including cancer^30–33^. As dysbiosis is frequent in stomach cancer^34, 35^, and considering the mechanism of action of leucovorin, it may be of interest to study whether organisms from these two genera may sequester or prevent the bacterial production of folinic acid^36^.

We found three microbial genera whose abundances were strongly associated with breast cancer response to doxetaxel. Indeed, the involvement of the microbiome in breast cancer (BRCA)^22, 37^ has recently received considerable attention. In BRCA, we found that the genus containing Epstein-Barr virus (EBV) was negatively associated with response to docetaxel, which is concordant with previous findings that EBV is associated with chemoresistance to docetaxel in gastric cancer^38^. Interestingly, Cyanobacteria were predictive features in several cancers in our study and we identified a genus of Cyanobacteria as predictive of response to docetaxel in BRCA. Notably, the presence of Cyanobacteria in BRCA was recently confirmed by Nejman et al.^22^ by 16S-rRNA sequencing. While the genus we identified, *Raphidiopsis*, a planktonic Cyanobacteria that produces toxins harmful to human health and found in freshwater, is possibly a taxonomic identification error in the original microbial abundance estimates, our findings may point to a related genus under the recently discovered clade Melainabacteria of Cyanobacteria^39^, which is present in humans. Though Melainabacteria are difficult to culture, we believe that confirmation of the relationship between BRCA to response to docetaxel and Melainabacteria should be tested, and a first step would be to confirm our computationally derived findings in a dedicated 16S-rRNA analysis.

Interestingly, in sarcoma (SARC), among the most predictive microbial features we found the genus *Lactococcus* to be positively associated with response to docetaxel. *Lactococcus* contains species that can sometimes cause opportunistic infections in humans, as *Lactococcus* are similar to *Streptococcus* and formerly belonged to that genus. The result that this genus was positively associated with response in our model initially appeared counterintuitive, although while the use of therapeutic bacteria as antitumor agents has not been an extensively studied field, there have been some limited findings in the literature that suggest the use of bacteriotherapy as anticancer agents^40^. Historically, the intentional use of the toxins of various *Streptococcus* species showing significant antitumor activity in SARC has been documented^41–43^. One possible testable explanation for some microbes being strongly positive predictive of docetaxel response in our model is that they might produce some extracellular products or toxins that could work as an adjuvant to the chemotherapy.

In summary, while these findings and others reported in this study are computationally derived associations, we believe that they can serve as leads for further experimental studies of the role of microbial species in modulating patients survival and drug response, potentially by metabolizing drug levels in the tumor microenvironment as suggested above, or by altering the immune response, either by changing the levels of specific immunometabolites or by having the tumors present specific bacterial antigens^44^.

## Methods

### Data retrieval and processing

Normalized and batch effect corrected microbial abundance data for 32 TCGA tumor types were downloaded from the online data repository referenced in Poore et al.^10^ (ftp://ftp.microbio.me/pub/cancer_microbiome_analysis). Specifically, the “Kraken-TCGA-Voom-SNM-Plate-Center-Filtering-Data.csv” microbial abundance data file and adjoining “Metadata-TCGA-Kraken-17625-Samples.csv” metadata file were used as the starting input for further data processing.

We first filtered the data for primary tumor samples (TCGA “Primary Tumor” or “Additional - New Primary” sample types). Poore et al. generated microbial abundances from all the available WGS and RNA-seq data in legacy TCGA (after some quality filters), which frequently contained replicate WGS and RNA-seq data for the same case and sample type. It was common in legacy TCGA to increase WGS sequencing coverage by performing an additional sequencing run from the same sample and these secondary runs typically had a much lower number of reads and coverage compared to their corresponding primary sequencing runs. When comparing the normalized and batch effect corrected read counts between these WGS runs, we found that microbial abundance data which came from lower coverage secondary runs could be substantially different from abundances derived from the larger primary sequencing runs. Therefore, we excluded microbial abundance data which came from secondary runs. In addition, legacy TCGA commonly contained data for the same samples analyzed using different computational pipeline versions. We excluded replicate microbial abundance data from older TCGA analysis pipeline versions if a replicate from a newer version existed. After the above filters, the Poore et al. data went from 17,625 samples and 10,183 unique cases to 12,111 samples and 9,812 unique cases (comprising of 1,944 WGS samples from 1,904 unique cases and 10,167 RNA-seq samples from 9,745 unique cases).

TCGA gender, age at diagnosis, and tumor stage demographic and clinical data and as well as primary tumor RNA-seq read count data for the 32 TCGA tumor types included in our study were obtained from the NCI Genomic Data Commons (GDC Data Release v29.0) using the R Bioconductor package GenomicDataCommons. TCGA GENCODE v22 gene annotations were obtained from the GDC data portal and Ensembl Gene v98 using the R package rtracklayer and R Bioconductor packages AnnotationHub and ensembldb. The downloaded GDC primary tumor cohort with RNA-seq read count data comprised of 9,735 samples from 9,680 unique cases. There were 68 cases at the GDC which had missing age of diagnosis but existing values in the Poore et al. data and we chose not to exclude these data and used the Poore et al. age of diagnosis values for these cases. TCGA curated survival phenotypic data^45^ were obtained from UCSC Xena. Cases which had both missing overall survival (OS) and progressive-free interval (PFI) outcome data were excluded from survival modeling.

TCGA curated drug response clinical data were compiled from Ding et al.^14^ Our drug response models used the following binary classification targets: complete response (CR) and partial response (PR) were labeled as responders and stable disease (SD) and progressive disease (PD) as non-responders. All TCGA samples with drug response phenotypic data were from pre-treatment biopsies. Due to the limited cancer-drug combination cohort sizes in TCGA, we modeled each drug individually, even if a patient received multiple drugs concurrently. If the same drug was given at multiple timepoints to a patient, we only considered their first drug response. We considered cancer-drug combinations that contained a minimum of 18 cases and at least 4 cases per response binary class, except for STAD oxaliplatin, where we allowed a minimum of 14 cases so that the gene expression dataset could be included. In total, we analyzed 30 cancer-drug combinations which had paired microbial abundance and gene expression data that met the above thresholds. Combined feature microbial abundance and gene expression datasets were created by joining data from each individual dataset which had matching TCGA sample UUIDs. For some TCGA cases, data existed from multiple different aliquots per sample or multiple technical runs per aliquot, therefore in these cases all combinations were joined at the sample UUID level. Cross-validation sampling probability weights as well as model and scoring sample weights were applied to account and adjust for any imbalance caused by the process.

### ML modeling

Machine learning (ML) models were built using the scikit-learn^46^ and scikit-survival libraries^47–49^. Custom extensions to scikit-learn and scikit-survival were developed to add new methods and functionalities required by this project. Survival models were built using Coxnet – regularized Cox regression with elastic net penalties^11^. Coxnet models controlled for gender, age at diagnosis, and tumor stage clinical prognostic covariates by including them as unpenalized features in the model (Coxnet penalty factor = 0). Drug response classification models were built using three different ML methods: 1) a variant of the linear support vector machine recursive feature elimination (SVM-RFE) algorithm^15^ that we developed with a number of additional features and better performance than the scikit-learn built-in version, 2) logistic regression (LGR) with elastic net^16^ (L1 + L2) penalties and embedded feature selection, and 3) LGR with an L2 penalty and limma^17^ (for tumor microbial and combination datasets) or edgeR^18, 19^ (for RNA-seq count datasets) differential abundance/expression feature scoring inside a k-best wrapper feature selection method around the learning algorithm. Limma differential abundance analysis was run inside the ML pipeline with default parameters except for fitting an intensity-dependent trend to the prior variances and running a robust empirical Bayes procedure (eBayes function parameters trend = TRUE and robust = TRUE). edgeR differential expression analysis was run inside the ML pipeline with default parameters except for enabling robust estimation of the negative binomial dispersion (calcDispersions function robust = TRUE) and robust estimation of the prior quasi-likelihood (QL) dispersion (glmQLFit function robust = TRUE). Both limma and edgeR methods scored and ranked features by differential abundance/expression p-value.

All three drug response ML methods unconditionally included the same three clinical covariates in the model as in the prognosis models by having them bypass feature selection in the ML pipeline, though in drug response models, clinical covariates were modeled as L2 penalized features. In SVM-RFE, clinical covariate features bypassed recursive feature elimination but were always included at each RFE recursive feature elimination model fitting step as well as final model refitting. To the best of our knowledge, no available comprehensive ML library in python or R currently provides an elastic net LGR algorithm with the functionality to specify features that can bypass embedded feature selection and be modeled with an L2 penalty (setting the R glmnet penalty factor, for example, does not provide this functionality as it is not a penalty factor per regularization term but a factor applied to the sum of both L1 and L2 terms). In order to develop this functionality for our study, our elastic net LGR model pipeline was designed as a two-level LGR, 1) an elastic net LGR and embedded feature selection on only microbial abundance or gene expression features with clinical covariates bypassing this step, followed by 2) an L2 penalized LGR on features selected by the elastic net LGR step and the clinical covariates. We know this design does not likely produce the exact same model settings and results of a single-level elastic net LGR algorithm with the functionality we needed, if such an implementation it existed, though we tested every drug response model through an ML pipeline with elastic net LGR and no clinical feature selection bypass and found that model predictive performance, feature coefficients and signs, and feature importance rankings were similar to our two-level ML pipeline setup.

Gender was one-hot encoded and tumor stage ordinal encoded by major stage. In the final cohort included in our prognosis and drug response models, 3363 out of 9708 tumor microbial abundance cases (34.64%) and 3244 out of 9484 gene expression cases (34.21%) had tumor stage “not reported” or BRCA stage “X”. Since missing tumor stage metadata is so prevalent in TCGA, we took the approach of including these in our study and modeled missing tumor stage with as neutral an ordinal encoding as possible. Looking at the distribution of reported major tumor stages in our cohort, we determined that encoding missing data as an ordinal between tumor stage II and III was as close to the middle of the distribution of stages in TCGA as we could possibly achieve with ordinal encoding.

All prognosis and drug response models included the previously described feature selection as well as normalization and transformation steps integrated into the ML modeling pipeline using an extended version of the scikit-learn Pipeline framework. Each cancer, data type, and survival or drug response target type combination was modeled individually using a nested cross-validation (CV) strategy to perform model selection and evaluation on held-out test data. Training data splits always underwent feature selection, normalization, and transformation through the ML pipeline independently from held-out test or validation data splits before learning. Models built using gene expression read count data included edgeR low count filtering, weighted trimmed mean of M-values (TMM) normalization, and log counts per million (CPM) transformation steps within the ML pipeline. These were developed and integrated into our scikit-learn-based framework via R and rpy2. All models also included standardization of features within the ML pipeline just before learning. During prediction, held-out test or validation data were feature selected, normalized, and transformed through the ML pipeline using the parameters learned from the training data at each pipeline step before model prediction and scoring. Hyperparameter search and optimization of all model pipeline steps was performed in nested fashion within the inner nested CV. All cross-validation iterators kept replicate sample data per case grouped together such that data would only reside in either the train or test split during each CV iteration.

Survival models used a stratified and randomly shuffled outer CV with 75% train and 25% test split sizes that was repeated 100 times. The CV procedure stratified the splits on event status. Each training set from the outer CV was used to perform hyperparameter tuning and model selection by optimizing Harrell’s concordance index (C-index) over a stratified, randomly shuffled, 4-fold inner CV on the training set repeated 5 times. A few cancer datasets contained fewer than four uncensored cases which required reducing the number of inner CV folds for these models such that at least one case per fold was uncensored. The data derived from Poore et al. often included more than one sample per case, and an unequal number of samples between cases, therefore requiring either ML model sample weighting or CV random sampling per case. The Coxnet implementation in scikit-survival does not currently support sample weights, therefore our custom outer CV iterator randomly sampled one replicate sample per case during each iteration, using a sampling procedure with probability weights that balanced the probability that a replicate WGS- or RNA-seq-based sample was selected during each CV iteration. Model selection grid search was performed on the following hyperparameters: elastic net penalty L1 ratios 0.1, 0.3, 0.5, 0.7, 0.8, 0.9, 0.95, 0.99, and 1, and for each L1 ratio a default alpha path of 100 alphas using an alpha min ratio of 10^−2^. Alpha is the constant multiplier of the penalty terms in the Coxnet objective function. Optimal alpha and L1 ratio settings were determined via inner CV and a model with these settings was then refit on the entire outer CV train data split. Model performance was evaluated in both inner and outer CV on each held-out validation or test data split, respectively, by generating model test predicted risk scores and using these scores to directly calculate C-index scores. We also evaluated and compared model predictive performance for each test data split survival time period by calculating time-dependent cumulative/dynamic AUCs^12, 13^.

Drug response models used a stratified, randomly shuffled, 4-fold outer CV that was repeated 25 times (i.e., 100 model instances). Each training set from the outer CV was used to perform hyperparameter tuning and model selection by optimizing the area under the receiver-operating characteristic curve (AUROC) over a stratified, randomly shuffled, 3-fold inner CV repeated 5 times. Case replicate sample weights were provided to SVM-RFE and LGR learning algorithms and all model selection and evaluation scoring methods. Class weights were provided to SVM-RFE and LGR learning algorithms to adjust for any class imbalance. Model selection grid search was performed on the following hyperparameters: L2 penalized SVM and LGR C regularization parameter from a range of 10^−5^ to 10^3^, elastic net LGR L1 ratios of 0.1, 0.3, 0.5, 0.7, 0.8, 0.9, 0.95, 0.99 and 1, elastic net LGR C regularization parameter from a range of 10^-2^ to 10^3^ (microbial abundance) or from 10^-2^ to 10^1^ (gene expression and combined data type), and finally RFE, elastic net LGR, and limma and edgeR feature scorer k-best feature selection search range from 1 to 400 top scoring microbial abundance, gene expression, or combined data type features. SVM-RFE models performed a feature elimination procedure of the one worst feature per recursive step for microbial abundance models (which started with 1287 features in the Poore et al. data) and 5% of worst remaining features per recursive step until 1300 features were reached followed by the one worst feature per recursive step for gene expression (starting with 60,483 features in GENCODE v22) and combined data type models (starting with 61,770 features). Optimized hyperparameter settings were determined via inner CV and a model with the optimized settings was then refit on the entire outer CV train data split. Model performance was evaluated in both inner and outer CV on each held-out validation or test data split, respectively, by AUROC, average precision (AVPRE) or area under precision-recall curve (AUPRC), and balanced accuracy (BCR). AUROC was used to evaluate and select the best model and optimized hyperparameter settings from the grid search.

Gender, age at diagnosis, and tumor stage clinical covariate-only survival models were built using standard unpenalized Cox regression. Clinical covariate-only drug response models were built using L2 penalized linear SVM or LGR. Models included standardization of features as part of the ML pipeline. Models were trained and tested using the same outer CV iterators and train/test data splits as their corresponding microbial abundance, gene expression, or combination data type models. To test whether a Coxnet, SVM-RFE, or LGR microbial abundance or gene expression model was significantly better than their corresponding Cox, linear SVM, or LGR clinical covariate-only model, respectively, a two-sided Wilcoxon signed-rank test was performed between the 100 pairs of C-index or AUROC scores between both models. All raw p-values generated from the signed-rank test across survival or drug response analyses from the same data type were adjusted for multiple testing using the Benjamini-Hochberg (BH) procedure to control the false discovery rate (FDR), and a threshold FDR ≤ 0.01 was used to determine statistical significance. To test whether a combined data type model was significantly better than its corresponding microbial abundance or gene expression model, a two-sided Dunn test was performed between all three groups of data type model scores. Each Dunn test raw p-value was adjusted for multiple testing using the Benjamini-Hochberg (BH) procedure to control the false discovery rate (FDR), and a threshold FDR ≤ 0.05 was used to determine statistical significance.

Permutation tests were performed by shuffling dataset class labels 1000 times and each time running the outer CV procedure on the permuted dataset, where for each CV iteration we fit a model instance and calculated an AUROC score, totaling 100,000 fits and scores for each model. Permutation mean AUROC scores were compared to the true mean AUROC score for the model and a p-value was calculated from the fraction of permutation mean scores that were greater than or equal to the true mean score. A p-value ≤ 0.05 was used to determine statistical significance. The Freedman-Draconis rule was used in permutation test histogram plots to compute the bin width. Analysis of the effect of number of selected features on model performance was performed via the hyperparameter grid search and tuning that occurred in the nested inner CV during each model instance fitting, where scores for every combination of hyperparameter setting and inner CV train/validation fold were saved for all model instances and used for plotting.

### Microbial abundance model feature analysis

For each analysis, 100 prognosis or drug response model instances were generated from the outer CV procedure. Each model instance selected a subset of features that performed best during CV and the model algorithm learned coefficients (or weights) for each feature. To select microbial genera for downstream investigation from the feature results across all these model instances, we proceeded as follows. First, we applied a two-sided Wilcoxon signed-rank test that the mean feature coefficient rank generated by the model is shifted away from zero, and thus that the genus is identifiably positively or negatively associated with survival or drug response. For all Wilcoxon tests, we used the package coin^50^, which allows exact calculation of p-values. Coefficients were ignored when a genus was assigned a zero coefficient or absent from a model. Second, within each model, all coefficients, ignoring the results of the Wilcoxon test, were ranked by absolute magnitude. We then kept genera that were among the top 50 features in at least 20% of the models and for which the Holm-adjusted, two-sided Wilcoxon signed-rank test p-value was ≤ 0.01. Having a Coxnet feature coefficient equal to zero or feature being absent from an SVM-RFE or LGR model was not strong enough evidence that the genus has no effect, but rather that one or more features with stronger effect were chosen. Thus, we ignored genera with a zero coefficient or absent from a model when computing mean coefficient weight and Wilcoxon statistics on the means.

For the drug response models, where three ML methods were tested, we noted the features selected by individual models and the median rank the feature attained in the instances in which it appeared, but further filtered the features to account for the consensus between ML models. We kept features selected in any two ML model methods that individually met our criteria for inclusion, ignoring features in ML models that did not meet these criteria. We then computed the Spearman correlation between the median ranks attained by the features.

For each selected microbial feature, we tested whether it was a significantly univariate feature of survival or drug response. This is a strictly different question than whether the coefficient of a feature has consistent sign – sign may be consistent when used in combination with other features, but the feature may not be individually predictive. For drug response models, we divided individuals into responders and non-responders, and for survival data we divided individuals whose survival time was greater or less than the censored median, ignoring those who were lost to follow up before median time. For each cancer-test type pair, we applied a two-sided Wilcoxon rank-sum test. We applied a Benjamini-Hochberg multiple hypothesis correction for each cancer-test type pair and report the false discovery rate in **Supplementary Data 2**.

We analyzed the distribution of features, selected by the rules described above, that had positive or negative signs for their mean coefficient. We used a two-sided binomial test to show that selected features had significantly more negative the positive mean coefficients. We used a two-sided Fisher’s exact test to determine if selected genera belonging to Firmicutes had a statistically significant difference in the breakdown between positive and negative mean coefficients than selected features as a whole.

## Data availability

All results generated from this work are available under https://doi.org/10.5281/zenodo.5221525.

## Code availability

All code and data used to produce this work are available under https://github.com/ruppinlab/tcga-microbiome-prediction.

## Supporting information

Description of Supplementary Data Files

## Acknowledgements

The results shown here are in whole or part based upon data generated by the TCGA Research Network (https://www.cancer.gov/tcga). This research was supported by the Intramural Research Program of the National Institutes of Health, National Cancer Institute and by NIH grant 1ZIABC011803-03. The authors would like to personally thank Christopher Buck from the NCI, Pedro Milanez-Almeida and John Tsang from NIH NIAID, and Alejandro Schäffer, Welles Robinson, Fiorella Schischlik, Sanju Sinha, and Sanna Madan from NCI CDSL for their assistance in this project. This study utilized the high-performance computational capabilities of the HPC Biowulf Linux cluster at the National Institutes of Health, Bethesda, MD (https://hpc.nih.gov). The authors would like to thank Richard Lehr, Tim Miller, Wolfgang Resch, and Steve Fellini for their assistance in running this analysis on the NIH HPC Biowulf cluster. The authors would also like to thank Joel Nothman, Andreas Mueller, and Adrin Jalali from the scikit-learn core development team and scikit-survival author Sebastian Pölsterl for their assistance with developing extensions to their libraries. Figure 1 embedded image credits (all CC 3.0, BSD 3-Clause, or Apache 2.0 license): scikit-learn.org, scikit-survival.readthedocs.io, eli5.readthedocs.io, gdc.cancer.gov, github.com/Bioconductor/BiocStickers, Fiorella Schischlik, wikipedia.org.

## Author information

### Contributions

L.C.H., E.M.G., and E.R. designed the study. L.C.H. and E.M.G. performed all computational analyses and results interpretation. L.C.H., E.M.G., and E.R. wrote the paper.

### Corresponding authors

Correspondence to Eytan Ruppin.

## Ethics declaration

### Competing interests

All authors declare that they have no competing interests.

## Supplementary Tables

**Supplementary Table 1.** Prognosis model results comparison to Milanez-Almeida et al.

|  | Milanez-Almeida |  | Hermida & Gertz |  |  |  |  |  |
| --- | --- | --- | --- | --- | --- | --- | --- | --- |
| Cancer | Tested | Hit | Tested | Hit | Common Hit | New Hit | M-A Tested New Hit | Missing M-A Hit |
| ACC | OS | OS | OS, PFI | OS, PFI | OS | PFI |  |  |
| BLCA | OS | OS | OS, PFI | OS, PFI | OS | PFI |  |  |
| BRCA | PFI |  | OS, PFI | PFI |  | PFI | PFI |  |
| CESC | OS |  | OS, PFI | OS, PFI |  | OS, PFI | OS |  |
| CHOL | *EXCL* |  | OS, PFI |  |  |  |  |  |
| COAD | OS |  | OS, PFI |  |  |  |  |  |
| DLBC | *EXCL* |  | OS, PFI |  |  |  |  |  |
| ESCA | OS |  | OS, PFI |  |  |  |  |  |
| GBM | OS |  | OS, PFI |  |  |  |  |  |
| HNSC | OS | OS | OS, PFI | OS | OS |  |  |  |
| KICH | *EXCL* |  | OS, PFI |  |  |  |  |  |
| KIRC | OS | OS | OS, PFI | OS, PFI | OS | PFI |  |  |
| KIRP | OS | OS | OS, PFI | OS, PFI | OS | PFI |  |  |
| LAML | OS | OS | *EXCL* |  |  |  |  | OS |
| LGG | PFI | PFI | OS, PFI | OS, PFI | PFI | OS |  |  |
| LIHC | OS | OS | OS, PFI | OS, PFI | OS | PFI |  |  |
| LUAD | OS | OS | OS, PFI | OS | OS |  |  |  |
| LUSC | OS |  | OS, PFI | PFI |  | PFI |  |  |
| MESO | OS | OS | OS, PFI | OS, PFI | OS | PFI |  |  |
| OV | OS |  | OS, PFI |  |  |  |  |  |
| PAAD | OS | OS | OS, PFI | PFI |  | PFI |  | OS |
| PCPG | EXCL |  | OS, PFI | OS |  | OS |  |  |
| PRAD | PFI | PFI | OS, PFI | PFI | PFI |  |  |  |
| READ | PFI |  | OS, PFI |  |  |  |  |  |
| SARC | OS |  | OS, PFI | OS, PFI |  | OS, PFI | OS |  |
| SKCM | EXCL |  | OS, PFI | OS, PFI |  | OS, PFI |  |  |
| STAD | OS |  | OS, PFI | PFI |  | PFI |  |  |
| TGCT | OS |  | OS, PFI |  |  |  |  |  |
| THCA | PFI |  | OS, PFI |  |  |  |  |  |
| THYM | PFI |  | OS, PFI | OS |  | OS |  |  |
| UCEC | OS |  | OS, PFI | OS, PFI |  | OS, PFI | OS |  |
| UCS | OS |  | OS, PFI |  |  |  |  |  |
| UVM | OS | OS | OS, PFI | OS, PFI | OS | PFI |  |  |

**Supplementary Table 2.** Per cancer, the number of times genera from the phylum Firmicutes were found among the selected features, whether positively or negatively associated with drug response or prognosis.

| Cancer | Selected Negatively | Selected Positively | Total |
| --- | --- | --- | --- |
| ACC | 8 | 13 | 21 |
| BRCA | 1 | 0 | 1 |
| CESC | 4 | 7 | 11 |
| ESCA | 0 | 4 | 4 |
| KIRC | 4 | 7 | 11 |
| LGG | 1 | 6 | 7 |
| SARC | 2 | 2 | 4 |
| STAD | 13 | 10 | 23 |

## Supplementary Figures

**Supplementary Figure 1.**
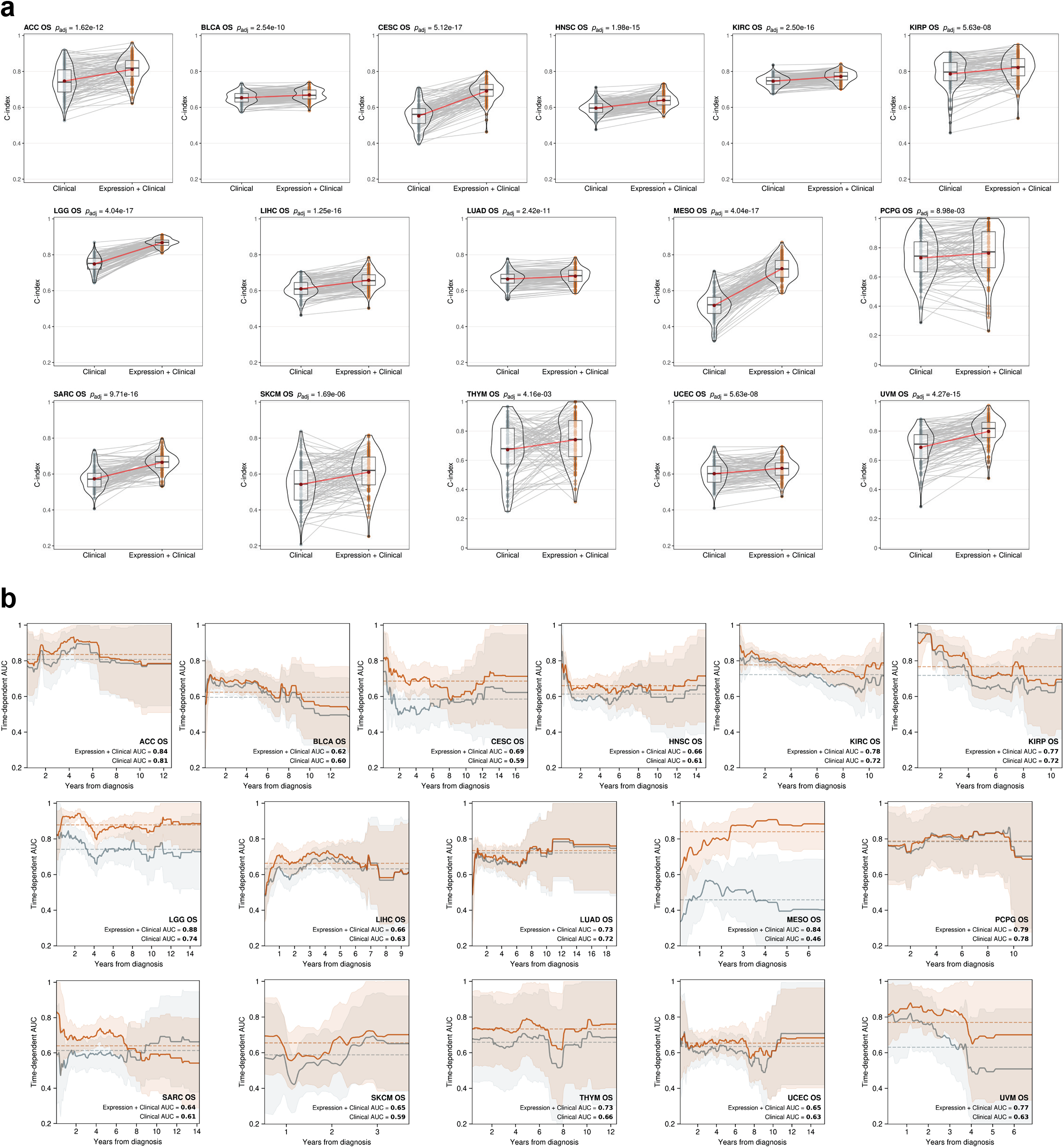
Performance of gene expression overall survival (OS) models in the 16 tumor types where gene expression adds predictive power to clinical covariates. **(a)** C-index score density distributions for gene expression with clinical covariate models vs clinical covariate-only models. Lines connecting points (light grey) represent score pairs from same train-test split on the data. Mean C-index scores and connecting lines shown in red. Significance for the prediction improvement over clinical covariate-only models was calculated using a two-sided Wilcoxon signed-rank test and adjusted for multiple testing using the Benjamini-Hochberg method with adjusted p-values shown at top. **(b)** Time-dependent, cumulative/dynamic AUCs for gene expression with clinical covariate models (orange) vs clinical covariate-only models (grey) following years after diagnosis. Mean AUCs across entire test time range after diagnosis shown as a horizontal dotted line and in legends and shaded areas denote standard deviations.

**Supplementary Figure 2.**
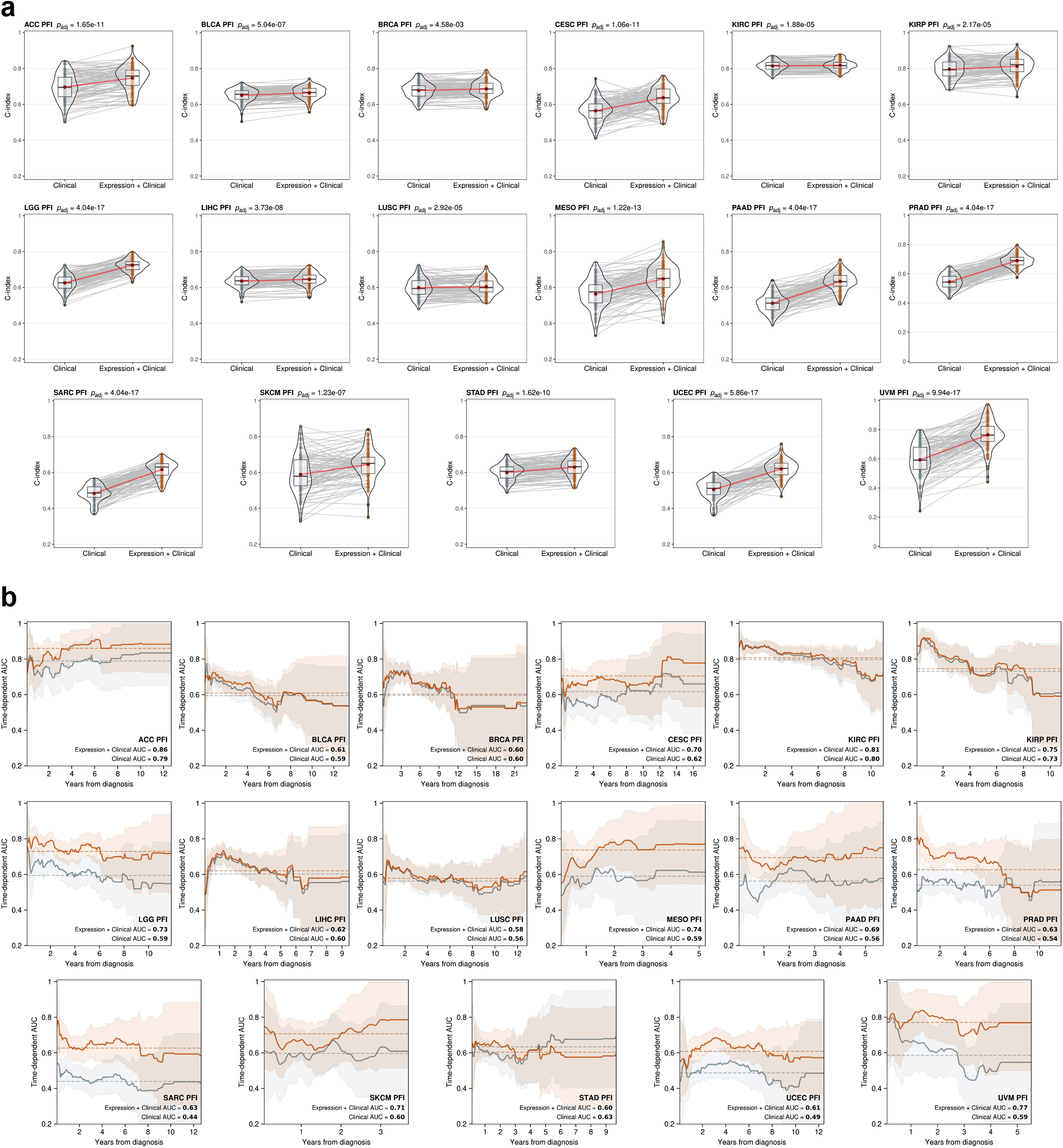
Performance of gene expression progression-free interval (PFI) models in the 17 tumor types where gene expression adds predictive power to clinical covariates. **(a)** C-index score density distributions for gene expression with clinical covariate models vs clinical covariate-only models. Lines connecting points (light grey) represent score pairs from same train-test split on the data. Mean C-index scores and connecting lines shown in red. Significance for the prediction improvement over clinical covariate-only models was calculated using a two-sided Wilcoxon signed-rank test and adjusted for multiple testing using the Benjamini-Hochberg method with adjusted p-values shown at top. **(b)** Time-dependent, cumulative/dynamic AUCs for gene expression with clinical covariate models (orange) vs clinical covariate-only models (grey) following years after diagnosis. Mean AUCs across entire test time range shown as a horizontal dotted line and in legends and shaded areas denote

**Supplementary Figure 3.**
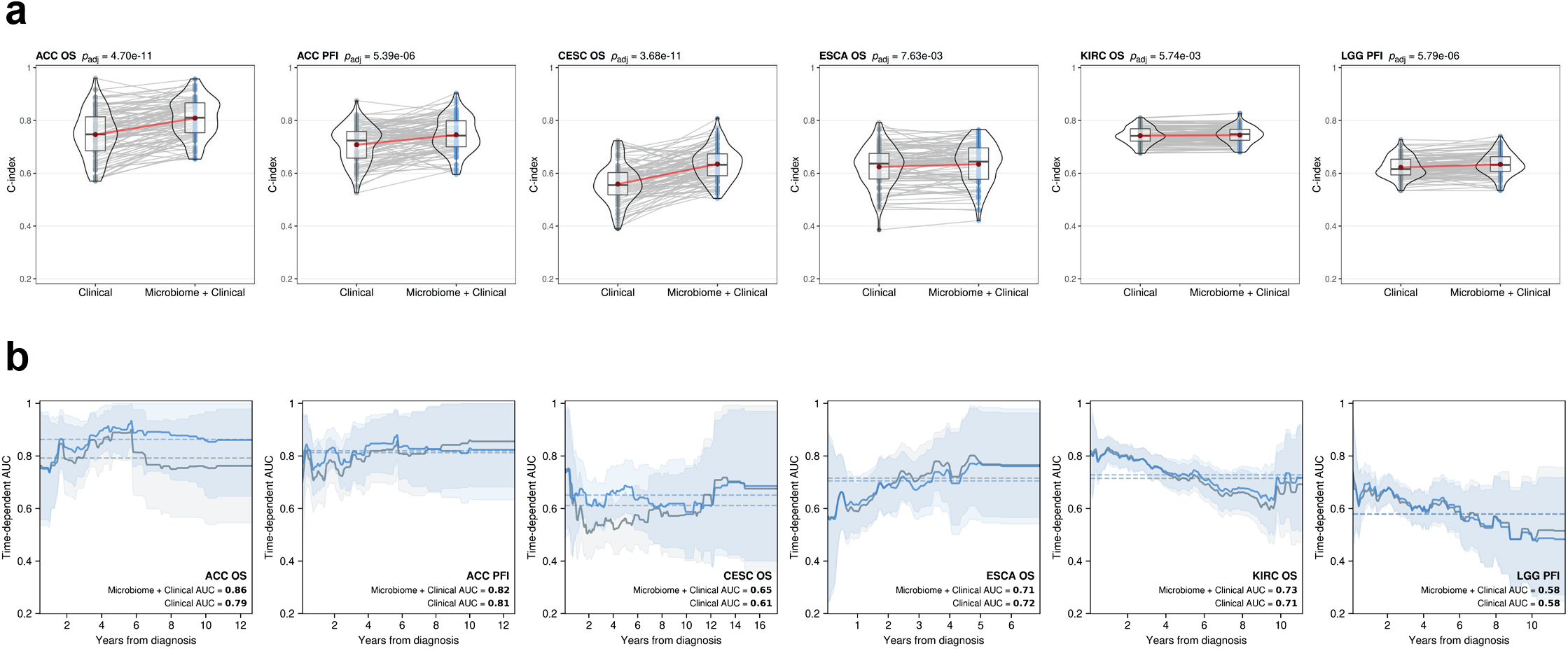
Performance of microbial abundance prognosis models in the five tumor types where microbial abundance features add predictive power to clinical covariates. **(a)** C-index score density distributions for microbial abundance with clinical covariate models vs clinical covariate-only models. Lines connecting points (light grey) represent score pairs from same train-test split on the data. Mean C-index scores and connecting lines shown in red. **(b)** Time-dependent, cumulative/dynamic AUCs for microbial abundance with clinical covariate models (blue) vs clinical covariate-only models (grey) following years after diagnosis. Mean AUCs across entire test time range shown as a horizontal dotted line and in legends and shaded areas denote standard deviations. Significance was calculated using a two-sided Wilcoxon signed-rank test and adjusted for multiple testing using the Benjamini-Hochberg method with adjusted p-values shown at top of violin plots in **(a)**.

**Supplementary Figure 4.**
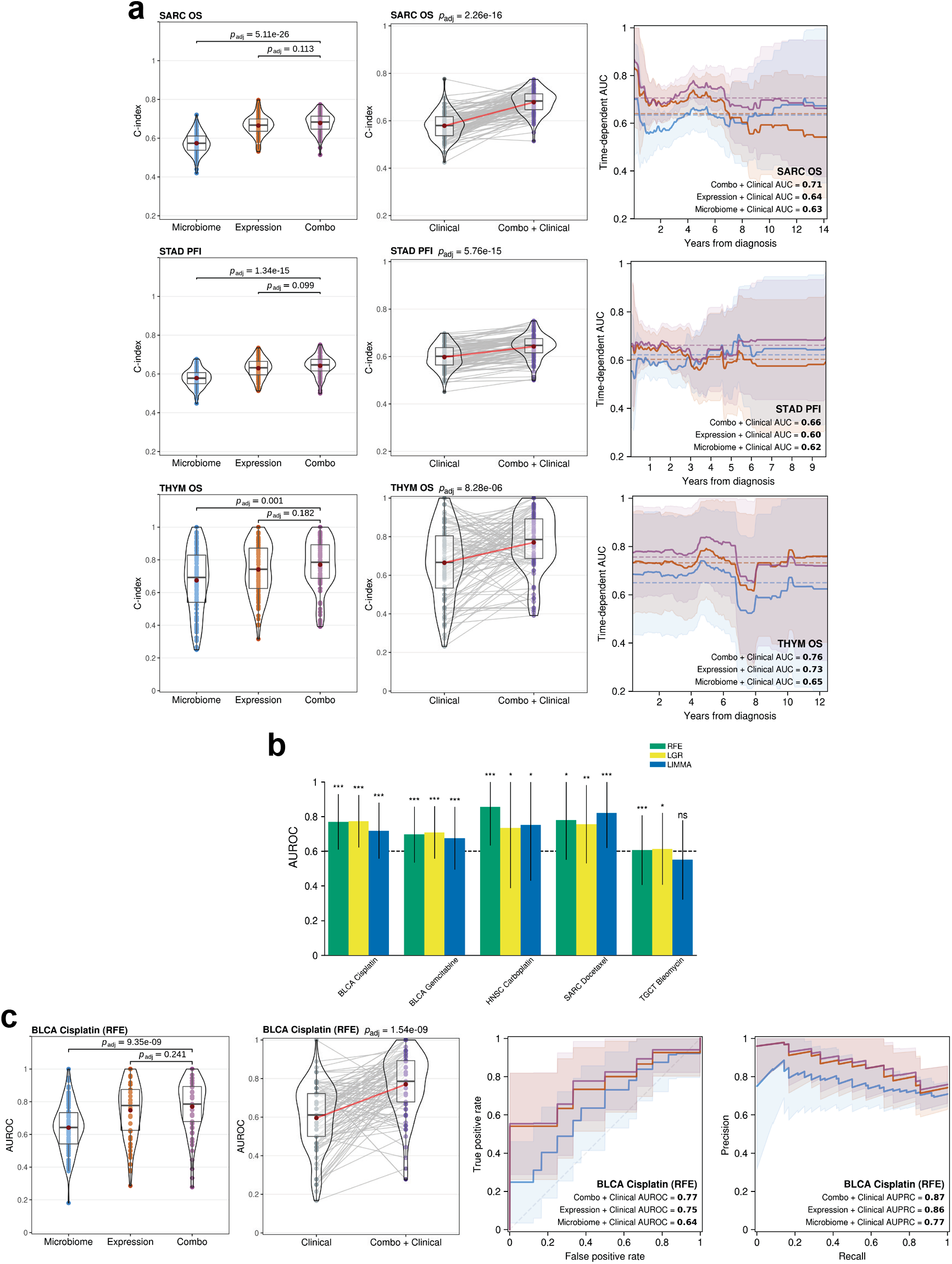
Performance of combined microbial abundance and gene expression models where combining both data types adds to predictive power. **(a)** Prognosis model C-index score violin density distributions (left) between microbial abundance, gene expression, and combined data type models. Mean scores shown in red. Significance was calculated using a two-sided Dunn test and p-values were adjusted for multiple testing using the Benjamini-Hochberg method. Model C-index score violin density distributions (middle) for combined data type with clinical covariate models vs clinical covariate-only models. Time-dependent, cumulative/dynamic AUCs (right) for combined data type (purple), microbial abundance (blue), and gene expression (orange) models following years after diagnosis. **(b)** ML method drug response model mean AUROC scores where combining data types performed better than clinical covariates alone. For the combined data type drug response model closest to reach significant improvement over respective single data type models **(c)** AUROC score violin density distributions (left) comparing each data type model, AUROC score violin density distributions (middle left) comparing combined data type and clinical covariate model to clinical covariate-only model, and mean ROC and PR curves (right) for each data type model. Mean AUROC and AUPRC scores shown in panel legends and shaded areas denote standard deviations.

**Supplementary Figure 5.**
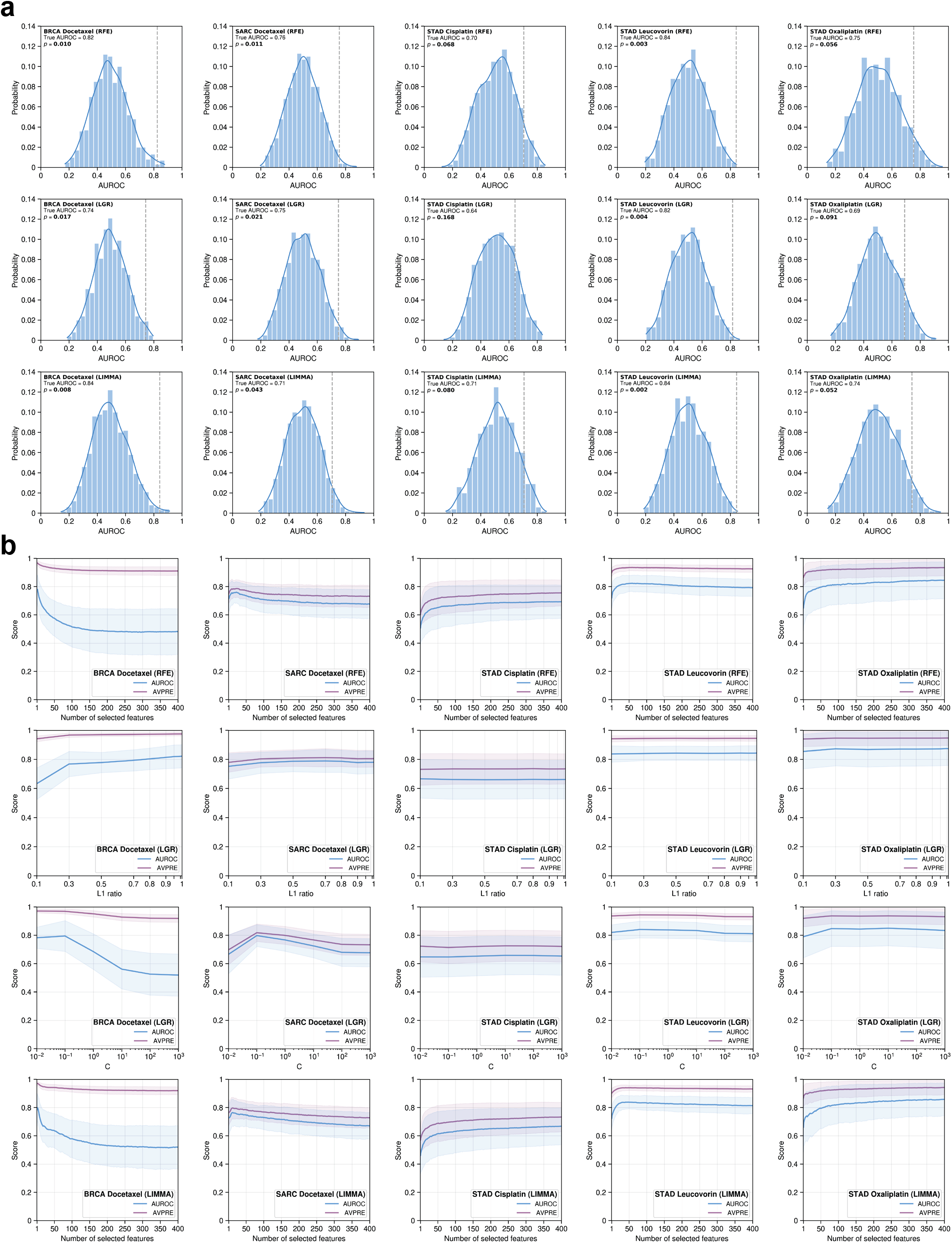
Evaluation of microbial abundance drug response model robustness. **(a)** Class label permutation test result histograms and significance showing the distribution of permutation mean AUROC scores. True mean AUROC score shown as dotted vertical grey line and kernel density estimate shown as a curve over the histogram. **(b)** Curves showing the effect that model hyperparameters which control the number of selected features had on mean AUROC and average precision (AVPRE) scores during hyperparameter grid search across all 100 model instances. Shaded areas denote standard deviations.

**Supplementary Figure 6.**
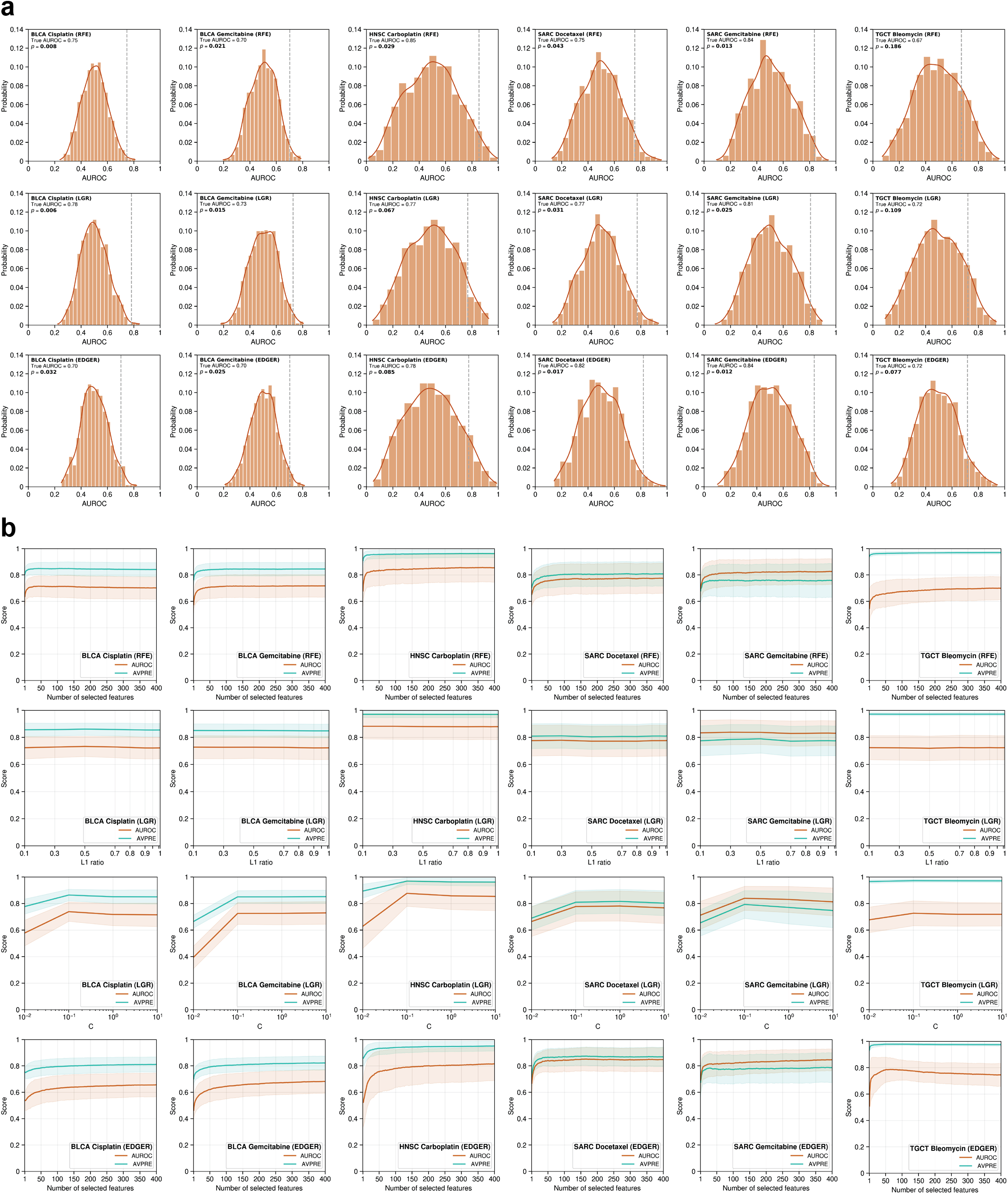
Evaluation of gene expression drug response model robustness. **(a)** Class label permutation test result histograms and significance showing the distribution of permutation mean AUROC scores. True mean AUROC score shown as dotted vertical grey line and kernel density estimate shown as a curve over the histogram. **(b)** Curves showing the effect that model hyperparameters which control the number of selected features had on mean AUROC and average precision (AVPRE) scores during hyperparameter grid search across all 100 model instances. Shaded areas denote standard deviations

## Notes

https://doi.org/10.5281/zenodo.5221525

https://doi.org/10.5281/zenodo.5838055

https://github.com/ruppinlab/tcga-microbiome-prediction

### Competing Interest Statement

The authors have declared no competing interest.

### Summary of Updates

Further revised and expanded manuscript and figures

## References

1. Grossman, R. L. et al. Toward a Shared Vision for Cancer Genomic Data. New England Journal of Medicine 375, 1109–1112 (2016).

2. Ahluwalia, P., Kolhe, R. & Gahlay, G. K. The clinical relevance of gene expression based prognostic signatures in colorectal cancer. Biochimica et Biophysica Acta (BBA) - Reviews on Cancer 1875, 188513 (2021).

3. Brodsky, A. S. et al. Expression profiling of primary and metastatic ovarian tumors reveals differences indicative of aggressive disease. PLoS One 9, e94476 (2014).

4. Liu, Y. et al. Pan-cancer analysis of clinical significance and associated molecular features of glycolysis. Bioengineered 12, 4233–4246 (2021).

5. Selfors, L. M., Stover, D. G., Harris, I. S., Brugge, J. S. & Coloff, J. L. Identification of cancer genes that are independent of dominant proliferation and lineage programs. Proc Natl Acad Sci U S A 114, E11276–E11284 (2017).

6. Shimoni, Y. Association between expression of random gene sets and survival is evident in multiple cancer types and may be explained by sub-classification. PLoS Comput Biol 14, e1006026 (2018).

7. Shukla, S. et al. Development of an RNA-Seq Based Prognostic Signature in Lung Adenocarcinoma. J Natl Cancer Inst 109, (2017).

8. Venet, D., Dumont, J. E. & Detours, V. Most random gene expression signatures are significantly associated with breast cancer outcome. PLoS Comput Biol 7, e1002240 (2011).

9. Milanez-Almeida, P., Martins, A. J., Germain, R. N. & Tsang, J. S. Cancer prognosis with shallow tumor RNA sequencing. Nature Medicine 26, 188–192 (2020).

10. Poore, G. D. et al. Microbiome analyses of blood and tissues suggest cancer diagnostic approach. Nature 579, 567–574 (2020).

11. Simon, N., Friedman, J., Hastie, T., & Tibshirani, R. Regularization Paths for Cox’s Proportional Hazards Model via Coordinate Descent. Journal of Statistical Software 39 (5), 1–13 (2011).

12. Hung, H. & Chiang, C.T. Estimation methods for time-dependent AUC models with survival data. Canadian Journal of Statistics 38 (1), 8–26 (2010).

13. Lambert, J. & Chevret, S. Summary measure of discrimination in survival models based on cumulative/dynamic time-dependent ROC curves. Statistical methods in medical research 25 (5), 2088–2102 (2016).

14. Ding Z et al. Evaluating the molecule-based prediction of clinical drug responses in cancer. Bioinformatics 32, (19): 2891–5 (2016).

15. Guyon, I., Weston, J., Barnhill, S. & Vapnik, V. Gene Selection for Cancer Classification using Support Vector Machines. Machine Learning 46, 389–422 (2002).

16. Zou H. & Hastie T. Regularization and variable selection via the elastic net. J R Statist Soc B 67 (2), 301–320 (2005).

17. Ritchie, M.E. et al. *limma* powers differential expression analyses for RNA-sequencing and microarray studies. Nucleic Acids Res 43 (7), e47 (2015).

18. Robinson, M. D., McCarthy, D. J. & Smyth, G. K. edgeR: a Bioconductor package for differential expression analysis of digital gene expression data. Bioinformatics 26, 139–140 (2010).

19. McCarthy, D. J., Chen, Y. & Smyth, G. K. Differential expression analysis of multifactor RNA-Seq experiments with respect to biological variation. Nucleic Acids Res 40, 4288–4297 (2012).

20. Geller, L. T. et al. Potential role of intratumor bacteria in mediating tumor resistance to the chemotherapeutic drug gemcitabine. Science 357, 1156–1160 (2017).

21. Pushalkar, S. et al. The Pancreatic Cancer Microbiome Promotes Oncogenesis by Induction of Innate and Adaptive Immune Suppression. Cancer Discov 8, 403–416 (2018).

22. Nejman, D. et al. The human tumor microbiome is composed of tumor type–specific intracellular bacteria. Science 368, 973–980 (2020).

23. Vabalas A. et al. Machine learning algorithm validation with a limited sample size. PLoS One 14 (11): e0224365 (2019).

24. Lin, D. et al. Microbiome factors in HPV-driven carcinogenesis and cancers. PLoS Pathog 16 (6), e1008524 (2020).

25. Zhu, H. et al. Chlamydia Trachomatis Infection-Associated Risk of Cervical Cancer: A Meta-Analysis. Medicine (Baltimore) 95 (13): e3077 (2016).

26. Gnanasekar A. et al. The intratumor microbiome predicts prognosis across gender and subtypes in papillary thyroid carcinoma. Comput Struct Biotechnol J 19, 1986–1997 (2021).

27. Dohlman et al. The cancer microbiome atlas: a pan-cancer comparative analysis to distinguish tissue-resident microbiota from contaminants. Cell Host Microbe 10 (2), 281–298.e5 (2021).

28. Parsonnet, J. et al. Helicobacter pylori infection and the risk of gastric carcinoma. N Engl J Med 325, 1127–1131 (1991).

29. Ferreira R.M. et al. Gastric microbial community profiling reveals a dysbiotic cancer-associated microbiota. Gut 67 (2), 226–236 (2018).

30. Abate, G., Qureshi, S. & Mazumder, S. A. Cedecea davisae bacteremia in a neutropenic patient with acute myeloid leukemia. J Infect 63, 83–85 (2011).

31. Akinosoglou, K. et al. Bacteraemia due to Cedecea davisae in a patient with sigmoid colon cancer: a case report and brief review of the literature. Diagn Microbiol Infect Dis 74, 303– 306 (2012).

32. Koh, Y. R. et al. The first Korean case of Sphingobacterium spiritivorum bacteremia in a patient with acute myeloid leukemia. Ann Lab Med 33, 283–287 (2013).

33. Wu, P. et al. Profiling the Urinary Microbiota in Male Patients with Bladder Cancer in China. Front Cell Infect Microbiol 8, 167 (2018).

34. Coker, O. O. et al. Mucosal microbiome dysbiosis in gastric carcinogenesis. Gut 67, 1024– 1032 (2018).

35. Castaño-Rodriguez N. et al. Dysbiosis of the microbiome in gastric carcinogenesis. Sci Rep 7 (1), 15957 (2015).

36. Ogwang, S. et al. Bacterial Conversion of Folinic Acid Is Required for Antifolate Resistance. Journal of Biological Chemistry 286, 15377–15390 (2011).

37. Eslami-S, Z., Majidzadeh-A, K., Halvaei, S., Babapirali, F. & Esmaeili, R. Microbiome and Breast Cancer: New Role for an Ancient Population. Front Oncol 10, (2020).

38. Shin, H. J., Kim, D. N. & Lee, S. K. Association between Epstein-Barr virus infection and chemoresistance to docetaxel in gastric carcinoma. Mol. Cells 32, 173–179 (2011).

39. Di Rienzi, S. C. et al. The human gut and groundwater harbor non-photosynthetic bacteria belonging to a new candidate phylum sibling to Cyanobacteria. eLife 2, e01102 (2013).

40. Sedighi M. et al. Therapeutic bacteria to combat cancer; current advances, challenges, and opportunities. Cancer Med 8 (6), 3167–3181 (2019).

41. Coley WB. Late results of the treatment of inoperable sarcoma by the mixed toxins of erysipelas and bacillus prodigiosus. Trans Southern Surg Gynecol Ass 18, 197 (1906).

42. Fehleisen F. Ueber die Züchtung der Erysipelkokken auf künstlichem Nährboden und ihre Übertragbarkeit auf den Menschen. Dtsch Med Wochenschr 8, 553-554 (1882).

43. Busch W. Aus der Sitzung der medicinischen Section vom 13 November 1867. Berl Klin Wochenschr 5, 137 (1868).

44. Kalaora S. et al. Identification of bacteria-derived HLA-bound peptides in melanoma. Nature 592 (7852), 138–143 (2021).

45. Liu, J. et al. An Integrated TCGA Pan-Cancer Clinical Data Resource to Drive High-Quality Survival Outcome Analytics. Cell 173, 400–416.e11 (2018).

46. Pedregosa et al. Scikit-learn: Machine Learning in Python. J Mach Learn Res 12, 2825–2830 (2011).

47. Pölsterl, S., Navab, N. & Katouzian, A. Fast Training of Support Vector Machines for Survival Analysis. in Machine Learning and Knowledge Discovery in Databases (eds. Appice, A., et al.) 243–259 (Springer International Publishing, 2015).

48. Pölsterl, S., Navab, N., & Katouzian, A., An Efficient Training Algorithm for Kernel Survival Support Vector Machines. 4th Workshop on Machine Learning in Life Sciences, 23 September 2016, Riva del Garda, Italy.

49. Pölsterl, S. et al. Heterogeneous ensembles for predicting survival of metastatic, castrate-resistant prostate cancer patients. F1000Res 5, 2676 (2017).

50. Hothorn, T., Hornik, K., Wiel, M. A. van de & Zeileis, A. Implementing a Class of Permutation Tests: The coin Package. Journal of Statistical Software 28, 1–23 (2008).

