## Supplementary material for "Predicting cancer prognosis and drug response from the tumor microbiome": Description of Supplementary Data Files

**Supplementary Data 1.xlsx**

**Worksheet #1 Prognosis model cohort sizes:** number of cases in each prognosis model dataset.

**Worksheet #2 Drug response model cohort sizes:** number of cases in each drug response model dataset, including the breakdown between responder (R) and non-responder (NR) cases.

**Supplementary Data 2.xlsx**

**Worksheet #1 Microbial genera features:** by cancer and by comparator, the genera selected as features by the criteria outline in Methods, and the ML methods that selected each genus. Further we show the number of times each genus was seen with rank at most 50 over the number of times the genus could be seen: 100, 200 or 300 if one, two, or three ML methods chose the feature, respectively. The column conditional direction shows the direction assigned to the feature for each ML method; an asterisk indicates that there was more than one relevant ML method, but all methods chose the same direction. The conditional direction is the typical sign of the coefficient within a model, which is conditional on other features in the model. Also shown is the Univariate FDR, which is the FDR that the feature is univariately predictive (n.s. is not significant), and the direction in which the feature is univariately predicative, which may differ from the conditional direction.

**Worksheet #2 Gene expression features**: by cancer and by comparator, the genes selected as features by the criteria outline in Methods, and the ML methods that selected each gene. Further we show the number of times each gene was seen with rank at most 50 over the number of times the gene could be seen: 100, 200 or 300 if one, two, or three ML methods chose the feature, respectively. The column conditional direction shows the direction assigned to the feature for each ML method; an asterisk indicates that there was more than one relevant ML method, but all methods chose the same direction. The conditional direction is the typical sign of the coefficient within a model.
